# Ablation of oligodendrogenesis in adult mice alters brain microstructure and activity independently of behavioural deficits

**DOI:** 10.1101/2023.07.06.547854

**Authors:** Malte S. Kaller, Alberto Lazari, Yingshi Feng, Annette van der Toorn, Sebastian Rühling, Christopher W. Thomas, Takahiro Shimizu, David Bannerman, Vladyslav Vyazovskiy, William D. Richardson, Cassandra Sampaio-Baptista, Heidi Johansen-Berg

## Abstract

Oligodendrocytes continue to differentiate from their precursor cells even in adulthood, a process that can be modulated by neuronal activity and experience. Yet, our understanding of the functional role of adult oligodendrogenesis remains limited. Previous work has indicated that conditional ablation of oligodendrogenesis in adult mice can lead to learning and memory deficits in a range of behavioural tasks. Our results, reported here, have replicated a key finding that learning to run on a complex wheel with unevenly spaced rungs is disrupted by ablation of oligodendrogenesis. However, using ex vivo MRI (MTR and DTI), we also found that ablating oligodendrogenesis by itself alters brain microstructure, independent of behavioural experience. Furthermore, in vivo EEG recording in behaviourally naïve mice with ablated oligodendrogenesis revealed altered brain activity in the form of increased EEG power density across a broad frequency range. Together, our data indicate that disrupting the formation of new oligodendrocytes directly alters brain microstructure and activity. This suggests a role for adult oligodendrogenesis in the maintenance of brain function and indicates that task-independent changes to brain structure and function might contribute to the learning and memory deficits associated with oligodendrogenesis ablation.

## Introduction

Myelination of axons plays a critical role in the functioning of the vertebrate nervous system by increasing the transmission speed and energy efficiency of neural processing (Nave, 2010; Salzer and Zalc, 2016). Recent studies have demonstrated that myelin is more dynamic than initially thought. Myelination can be stimulated by artificially exciting neuronal activity (Cullen et al., 2021; Gibson et al., 2014; Mitew et al., 2018), indicating that myelin plasticity, in addition to synaptic modification, might be one way in which experience can shape brain structure and function (Bonetto et al., 2021; Kaller et al., 2017; Xin and Chan, 2020). Indeed, changes in myelination and white matter (WM) microstructure have been consistently associated with learning in humans (Lakhani et al., 2016; Scholz et al., 2009) and rodents (Bacmeister et al., 2022; Sampaio-Baptista et al., 2013). In addition, adaptive myelination is proposed to regulate homeostatic coordination and oscillatory self-organization in local and large-scale brain networks (Dubey et al., 2022; Noori et al., 2020; Pajevic et al., 2022, 2014; Talidou et al., 2022). Thus, deficient myelin plasticity might lead to alterations in myelination and neural network function that impair neurological function (Geraghty et al., 2019; Knowles et al., 2022). Yet, the questions of when, how, and to what extent adaptive myelination contributes to relevant changes in neural circuit function require further investigation.

Myelination can be adaptively modified through the formation of new oligodendrocytes (OLs), the myelin forming glial cells in the central nervous system (Bergles and Richardson, 2016; Foster et al., 2019). OLs continue to differentiate even in adulthood (Hill et al., 2018; Hughes et al., 2018; Rivers et al., 2008; Young et al., 2013) from populations of oligodendrocyte precursor cells (OPCs) that remain abundant, dynamic and widespread in the CNS throughout life (Dawson et al., 2003). Recently, multi-photon imaging studies in live behaving mice have demonstrated continuous formation of oligodendrocytes and changes in oligodendroglial cells dynamics in response to motor learning (Bacmeister et al., 2022; Hill et al., 2018; Hughes et al., 2018). However, the contribution of such continuous and adaptive oligodendrocyte lineage dynamics to the functioning of the nervous system is still not well understood.

Transgenic mouse lines that allow conditional ablation of oligodendrogenesis in early adulthood have provided information about the causal involvement of new oligodendrocyte formation in behaviour. The first use of this transgenic approach revealed that mice with ablated oligodendrogenesis have an impaired ability to learn to run at speed on a complex wheel, suggesting a deficit in motor skills learning (McKenzie et al., 2014; Xiao et al., 2016). Subsequent studies found that interrupting OL generation also leads to deficits in long-term consolidation of spatial and fearful memories (Pan et al., 2020; Steadman et al., 2020). Additionally, reduced opioid reward learning (Yalçın et al., 2022) and impaired training-induced improvements in spatial working memory have also been reported (Shimizu et al., 2023). However, if and how the disruption of oligodendrogenesis in adulthood affects the CNS, independent of experience in a specific task, remains understudied.

The current study set out to further probe our understanding of behavioural, anatomical and physiological consequences of disrupting the formation of new OLs during adulthood. Our first aim was to replicate key findings indicating a motor learning deficit in adult mice with ablated oligodendrogenesis on the complex wheel task (McKenzie et al., 2014; Xiao et al., 2016). Our results replicated the finding that ability to perform a complex wheel (CW) running task is perturbed within the first hours of testing in animals with disrupted oligodendrocyte. Our second aim was to study how genetic ablation of OL differentiation affects brain microstructure and activity in adult mice using *ex vivo* magnetic resonance imaging (MRI) and *in vivo* EEG recording. We hypothesised that disruption of new oligodendrocyte formation alters the metrics of brain microstructure sensitive to myelin and, as a consequence, might modify cortical network activity. We investigated how learning to run the complex wheel affects brain microstructure in the presence and absence of ongoing OL differentiation, probing the mechanism by which adaptive OL formation might contribute to skill acquisition. We found alterations in both the microstructure and activity of the brain in animals with disruptioned oligodendrogenesis, irrespective of animal’s experience with the CW task.

## Results

### Deleting Myrf in OPCs leads to ablation of oligodendrogenesis

In the current study, we used a transgenic mouse model that allowed conditional deletion of myelin regulatory factor (*Myrf*) in the resident PDGFRα-positive OPC population (*Pdgfra-CreERT2:R26R-YFP: Myrf^(fl/fl)^*; referred to as ‘P-Myrf’ from here on; McKenzie et al., 2014). As *Myrf* encodes a transcription factor required for oligodendrocyte differentiation (Emery et al., 2009), animals with both alleles of *Myrf* conditionally deleted during adulthood (P-Myrf^(-/-)^) have markedly reduced formation of new oligodendrocytes 6 weeks after tamoxifen treatment, as compared to control animals with one functional allele (P-Myrf^(+/-)^) (Fig. 1A-D).

**Fig. 1.**
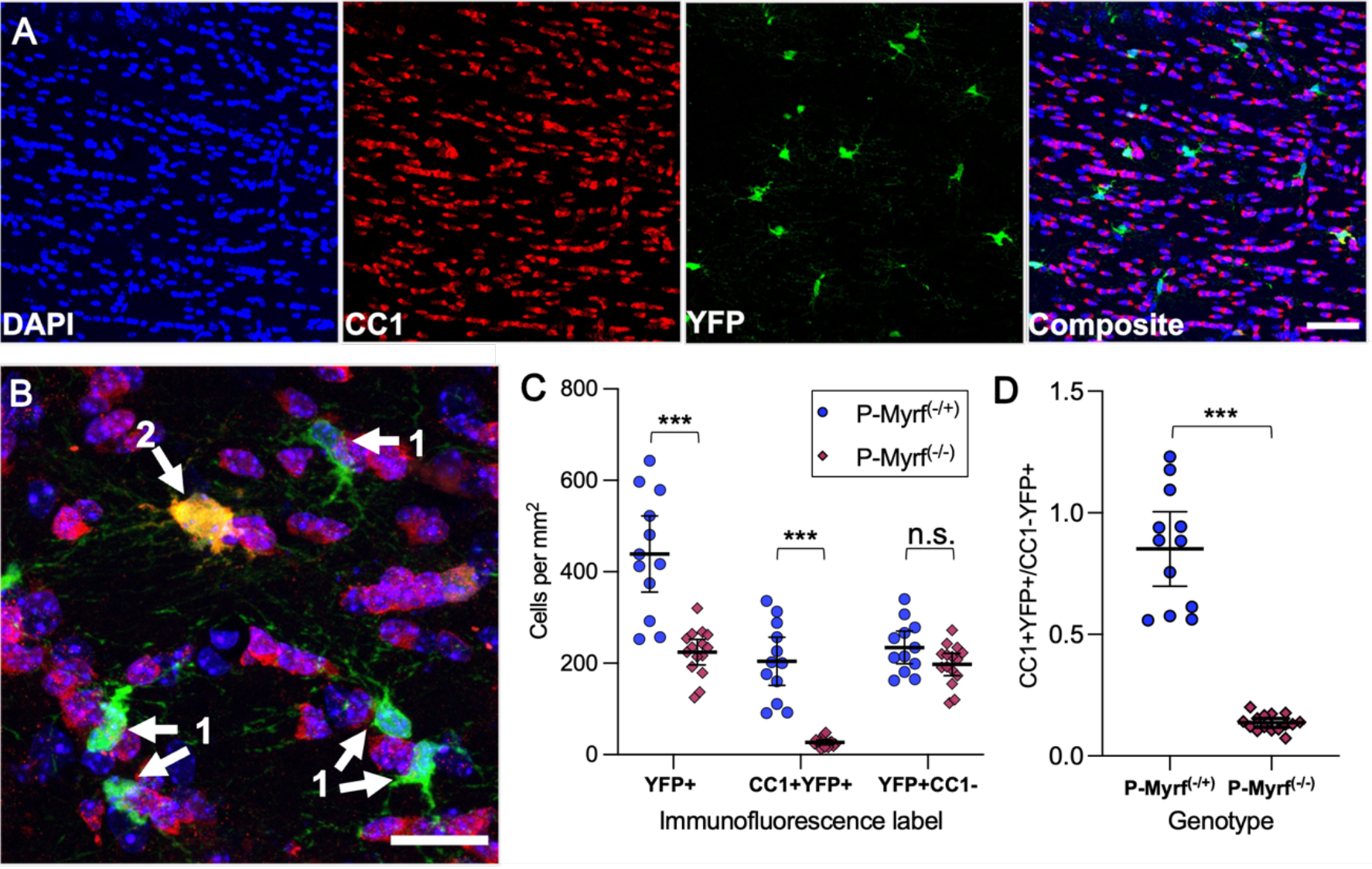
Deleting Myrf in OPCs significantly reduced the formation of new oligodendrocytes. **(A)** Representative individual image of immunohistochemistry, split into its separate channels. DAPI was used to identify individual cell nuclei, CC1 was used to label mature oligodendrocytes, and YFP marked OPCs at the time of tamoxifen administration (Scale bar 50 um). **(B)** CC1+ and YFP+ cells were quantified and labelled as either (1) an OPC that remained YFP+|CC1-, or (2) a recently matured Oligodendrocyte that was YFP+|CC1+. **(C)** Quantification confirmed a significant reduction of the density of YFP+|CC1+ recently matured oligodendrocytes in the P-Myrf(-/-) animals 6 weeks after tamoxifen administration. **(D)** The ratio between recently matured Oligodendrocytes (YFP+|CC1+) and labelled OPCs (YFP+|CC1-) was significantly lower in the P-Myrf(-/-). Data presented as Mean±95%CI, Welch’s two sample t-test used for statistical comparison. *** represents statistical significance of p<0.001. P-Myrf(+/-), n=12; P-Myrf(-/-), n=15.

### Ablation of oligodendrogenesis leads to impaired performance on the complex wheel

Previous work has indicated that such conditional ablation of oligodendrogenesis in adult mice leads to learning deficits in a range of different behavioural tasks (McKenzie et al., 2014; Pan et al., 2020; Steadman et al., 2020). We aimed to re-examine the ability of such mice to learn to run on a complex running wheel with irregularly spaced rungs (McKenzie et al., 2014; Xiao et al., 2016) (Illustrated in Fig. 2A).

**Fig. 2.**
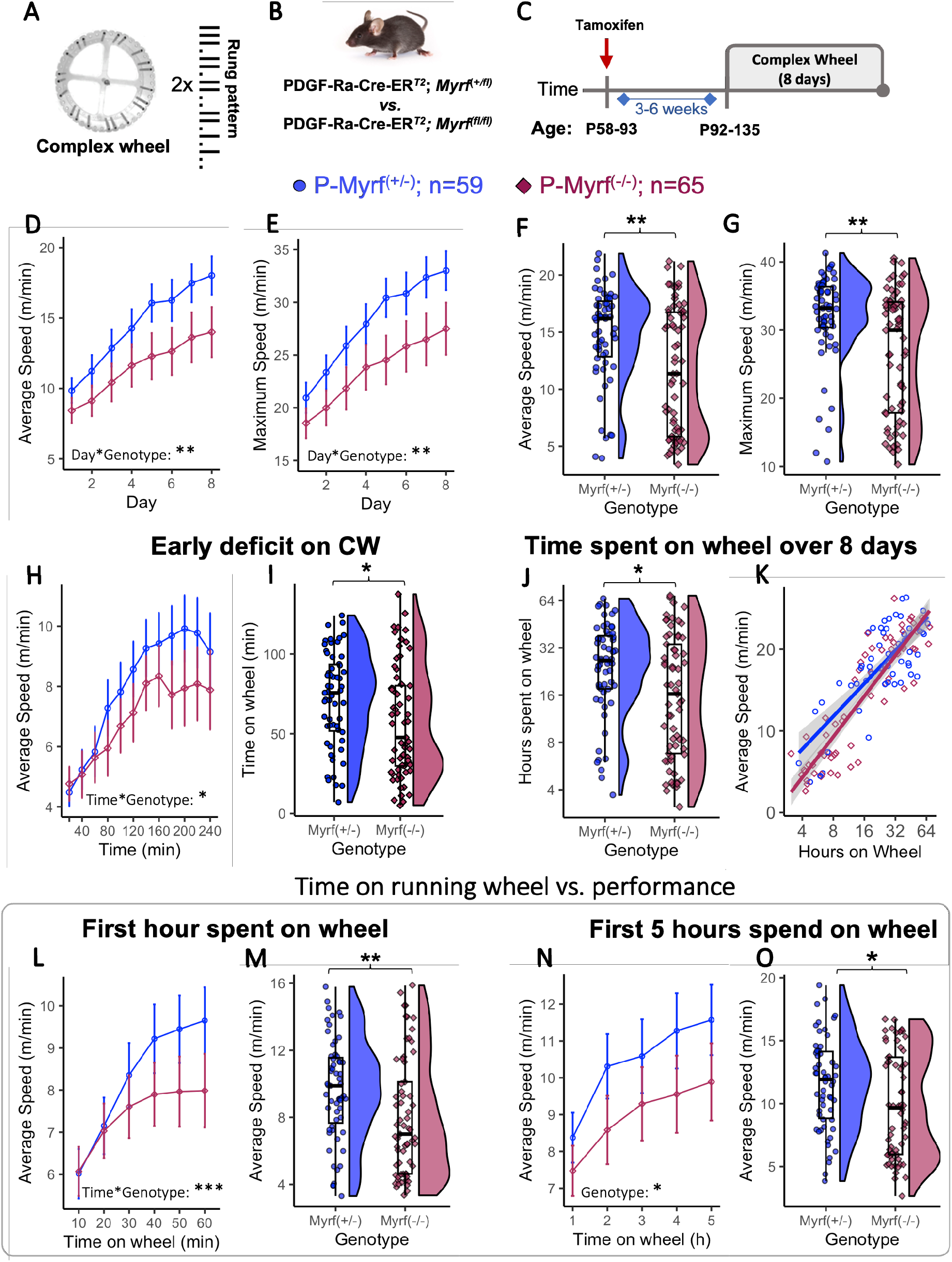
Preventing differentiation of new OL leads to impaired performance on the complex wheel. **(A)** Complex running wheel, with an irregular rung pattern displayed beside the wheel**. (B)** Animals bred on a PDGF-Rα-Cre-ERT2 back-ground with two floxed alleles of Myrf were compared to control animals that only have a single allele floxed. **(C)** Schematic Illustration of experimental design. Between 3-6 weeks after tamoxifen treatment, animals were given free access to the CW for 8 days. **(D)** Mean average speed and **(E)** maximum speed on the complex wheel of P-Myrf(-/-) was lower across 8 days. A statistically significant effect of Genotype and for Genotype*time interaction was detected for both speed metrics. **(F, G)** Raincloud plots of Average **(F)** and Maximum **(G)** speed across the 8 days of running. **(H)** Average running speed on the CW during the first 4 hours. P-Myrf(-/-) had a significantly lower speed improvement over the first 4 hours (Time*Genotype interaction). (**I)** Time spent on the wheel within the first 4 hours showed large variability and was lower within the P-Myrf(-/-) group (Mann-Whitney U, p=0.013). **(J)** P-Myrf(-/-) spent less time running on the complex wheel than their P-Myrf(+/-) siblings over 8 days of testing. **(K)** Statistically significant log relationship between the average speed of animals on day 8 of testing and the time animals spend on the wheel during the 8 days. **(L)** Average running speed plotted against the first hour of time animals spent on the wheel. **(M)** Average speed within 10 mins time period of wheel running after the first hour on the wheel. **(N)** Average running speed plotted against the time animals spend on the wheel for the first 5 hours (P-Myrf(+/-); n=58 ; P-Myrf(-/-); n=64). **(O)** Average speed within 1 hour of wheel running after 5 hours on the wheel. Data presented as Mean±95%CI or boxplots. Mixed-ANOVA (Main Factors: Genotype, Sex; Factor: Time) used for statistical comparison of performance over time comparison. Mann-Whitney U test used for statistical comparison of two groups. Asterix indicate statistical significance (*p<0.05, **p<0.01, ***p<0.001).

A total of 124 animals (P-*Myrf*^(+/-)^; n=59, 28 females; P-*Myrf*^(-/-)^; n=65, 29 females) were tested on the complex wheel (CW) across several experiments (see Methods). Animals received tamoxifen in early adulthood (age range: P60-90) and behavioural performance was tested 3-6 weeks later (Fig. 2C).

To investigate whether ablation of *Myrf* in OPCs during adulthood impacts performance on the complex running wheel, we tested for main effects of genotype, time and interactions on daily average and maximum speed using a mixed-two way ANOVAs. Sex was included as an additional between-subject factor.

Supplementary Table 2 contains comprehensive information on the statistical analysis methods utilized and the corresponding results for each dataset.

A main effect of genotype for average speed (F(1,120)= 10.00, *p=*0.002, Fig. 2D) and maximum speed ((F(1,120)=11.92, p<0.001, Fig. 2E) over 8 days confirmed that overall performance metrics of P-*Myrf*^(−/−)^ animals were lower than for their P-*Myrf*^(+/−)^ siblings. Furthermore, P-*Myrf*^(−/−)^ animals had impaired capacity to improve their performance with practice over training days (mixed two-way ANOVA, time*genotype interaction for average daily speed (F(2.74,328.5)=5.56, *p*=0.001, Fig. 2D) and maximum daily speed F(2.57,308.9)=4.76, *p*=0.005, Fig. 2E). Females outperformed males in terms of running performance (main effect of sex, average speed: F(1,120)= 13.1, p<0.001, maximum speed: F(1,120)= 17.5, p<0.001, Suppl. Fig. 1G) and spent more time on the complex running wheel (Suppl. Fig. 1H, Mann-Whitney U, *p*=0.014), yet no statistically significant interaction between animals’ sex and genotype was detected in any ANOVA test presented. Running ability on a normal running wheel and general motor skills, as tested by RotaRod and Balance Beam, were not impaired in the P-Myrf^(−/−)^ animals (Suppl. Fig. 1A-F), indicating that performance differences between genotypes did not generalize to other running tasks or tests of motor skill.

The average running speed (Fig. 2F) and maximum running speed (Fig. 2G) across the 8 days of training was lower in P-*Myrf*^(−/−)^ (Mann-Whitney U: average speed, Fig. 2F, *p*=0.003; maximum speed, Fig. 2G, *p*=0.002), further indicating a deficit in performance on the CW in animals with ablation of *Myrf* in OPCs during adulthood. However, the distribution of performance values was not equal between genotype groups, with higher variability and a bimodally shaped distribution especially prominent in the group of P-*Myrf*^(−/−)^ animals (Fig. 2F & G), a feature that was also reported by McKenzie et al. (2014).

Furthermore, the observed deficit on the CW was already detectable within the first 4 hours of exposure to the wheel (Mixed ANOVA: Genotype*Time F(4.83, 579.6) = 2.63, *p*=0.025, Fig. 2H), replicating an early deficit in complex wheel learning previously reported by Xiao et al (2016).

### P-Myrf^(−/−)^ animals spend less time running on the complex wheel

As running on the wheel was voluntary and animals were given unrestricted access to the wheels, the time individual animals spent on the running wheel varied significantly. Indeed, within the first 4 hours of exposure to the wheel, the time animals spent rotating the wheel ranged from about 5 minutes to over 2 hours (Fig. 2I). Interestingly, P-*Myrf*^(−/−)^ animals spent significantly less time on the wheel within this 4 hour period (Fig. 2I, Mann-Whitney U, *p*=0.013). A similar trend was observed over the 8 days of testing. Time running on the wheel ranged from around 4 hours to about 64 hours (Fig. 2H) and P-*Myrf*^(−/−)^ animals spent less time running on the complex running wheel than P-*Myrf*^(−/+)^ (Mann-Whitney U: *p*=0.014, Fig. 2H, also see Suppl. Fig. 1I). Similar to other performance metrics across 8 days (Fig. 2F & G), the time animals spent on the wheel had a wider and bimodal distribution in the P-*Myrf*^(−/−)^ group (Fig. 2H).

Importantly, we found a statistically significant log-positive relationship between the time animals spent running on the wheel and their average running speed on day 8 of testing (Fig. 2I, Spearman’s rank correlation, across genotype, rho = 0.800, p<0.001; P-*Myrf*^(+/−)^, rho=0.627, p<0.001; P-*Myrf*^(-/−)^, rho=0.871 p<0.001). This suggests that when considering running speed metrics over chronological time periods (e.g. days, Fig. 2D & E) it might be important to account for variation in the time animals spent running on the wheel, and therefore inter-individual variation in the time available for learning the task.

### Deficit on complex running wheel detected within the first hours of running

One way to account for such variation is to compare running performance of animals against the time they spent on the wheel, rather than chronological time. Hence, we analysed average running speed on the complex wheel for the first hour animals spent running on the wheel, regardless of when that running happened (Fig. 2 L & M). Using a mixed two-way ANOVA (main factors: genotype, sex; within subject factor: time(10 min intervals)), we found a significant effect of time (F(2.21, 265.3)= 88.0, p<0.001) and a significant time*genotype interaction for average daily speed (F(2.21, 265.3)= 9.06, p<0.001, Fig. 2J), indicating that P-*Myrf*^(−/−)^ animals showed less improvement in performance with practice over the first hour of running. The average running speed achieved in a 10-minute interval after 1 hour on the wheel was significantly lower in P-*Myrf*^(−/−)^ animals (Fig. 2K, Mann-Whitney U: *p*= 0.004).

As this deficit is prominent within the first hour spent running on the complex wheel, we wondered how the genotype differences changed over subsequent time animals spent on the wheel. Comparing performance across the first 5 hours running on the wheel, P-*Myrf*^(−/−)^ animals had a persistent lower average running speed than P-*Myrf*^(+/−)^ controls (Fig 2 L-O). A mixed two-way ANOVA (main factors: genotype, sex; within subject factor: time) confirms a statistically significant effect of time (average speed: F(2.2,259.9)= 53.8, p<0.001) and for genotype (average speed: F(1,118)= 4.99, *p*=0.027, Fig. 2N), yet not for an interaction between time spent on the wheel and the animals’ genotype that might indicate a deficit in learning ability over this time period (average speed: F(2.2,259.3)= 1.22, *p*=0.29). The running speeds in the 5^th^ hour of running on the wheel was different between genotypes (average speed: Mann-Whitney U: *p*= 0.031, Fig. 2O). This suggests the group differences that became apparent within the first hour on the wheel (Fig. 2L&M) persisted over subsequent hours of running.

Overall, these results replicate the main behavioural results obtained by McKenzie et al (2014) and Xiao et al (2016), indicating an early deficit in learning on the CW in animals with ablation of *Myrf* in OPCs during adulthood. However, unlike McKenzie et al. (2014), we found wide variation in the amount of time animals spent engaging with the task, a factor that is important to take into account when interpreting behavioural changes. Taking this variation into account by considering time spent on wheel rather than chronological time, we found that ablation of oligodendrogenesis results in deficits in performance improvement over the first hour of experience on the wheel. These deficits are maintained, but not exacerbated, over subsequent hours of experience.

### Probing task demands of the complex wheel task

To better understand the task demands of the wheel running task, we tested wild type (WT) mice on different variants of the wheel. Switching to a complex wheel after running on the normal wheel leads to a partial reduction in running speed in WT mice (Suppl. Fig. 5), that recovers after a day or two of further training, indicating the task demand specific to the CW is mainly probed in the first couple of days of exposure to the wheel.

To test whether mice learn a specific sequence of movements to master the complex wheel, we tested the effects of changing the rung sequence after exposure to one variant of the complex wheel. This does not lead to reduction in running speed in WT mice (Suppl. Fig. 6), suggesting that mice learn a general strategy for running on irregularly spaced rungs, rather than a specific sequence of movements.

Taken together, these behavioural findings in WT mice suggest that learning to run at speed on a complex wheel primarily involves adaptation to the irregular spaced rung positions, particularly in the early days of testing.

### Ablation of oligodendrogenesis alters brain microstructure independently of wheel running

Previous work has reported alterations in white matter microstructure, measured with MRI techniques, in response to motor learning (Sampaio-Baptista et al., 2020, 2013). However, the underlying mechanism is unclear, as these changes could relate to remodelling of pre-existing myelin and/or recruitment of new oligodendrocytes. Furthermore, the contribution of new oligodendrocytes to MRI metrics has not been directly tested.

In the current study we hypothesised that disruption of oligodendrogenesis in early adulthood affects brain microstructure, as it interferes with ongoing cellular dynamics and *de novo* myelination. Further, we hypothesised that learning to run on the CW affects brain microstructure. Finally, given evidence that adaptive oligodendrogenesis contributes to learning to run on the CW, we hypothesised that effects of CW running on brain microstructure should differ between knock-outs and controls.

We employed *ex vivo* MRI (see Methods) to compare P-Myrf^(−/−)^ (n=26) and P-Myrf^(−/+)^ (n=25) animals that had free access for 12 days to either the complex wheel (CW, n=26) or to a fixed immovable CW in which running was not possible (from here on referred to as ‘fixed wheel’, FW, n=25, Fig. 3A). This FW control condition was used to keep all environmental factors, other than wheel running, similar between groups (schematic illustration of experimental design, Fig. 3B).

**Fig. 3.**
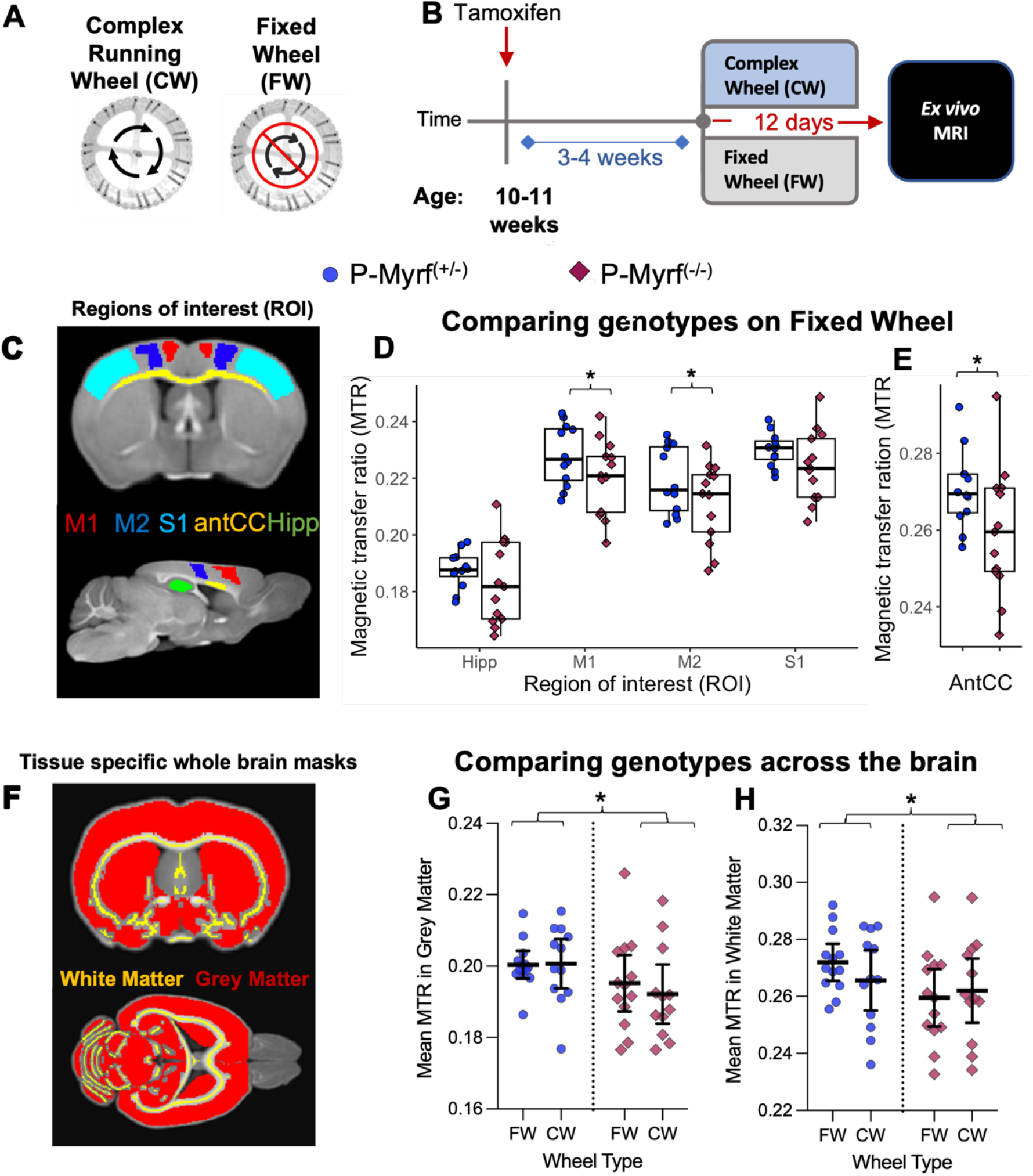
Ablation of oligodendrogenesis alters brain microstructure even in the absence of wheel running. **(A)** Animals used for ex vivo MRI imaging were exposed to either a complex wheel on which they can run, or an immovable complex wheel (Fixed Wheel, FW). **(B)** Illustration of experimental design. **(C)** Coronal and sagittal section of MTR image illustrating Regions of Interest (ROI) masks used for the ROI analysis. **(D)** For animals only exposed to the Fixed Wheel (FW, P-Myrf(+/-), n=12,, P-Myrf(-/-), n=13), ROI-specific mean MTR revealed that the ablation of oligodenrogenesis in the absence of wheel running lead to statistically significant changes in brain microstructure in the Primary motor cortex (M1), Secondary motor cortex (M2) and the (**E**) anterior corpus callosum (antCC). **(F)** Mask for the Grey Matter (GM, red) and White Matter skeleton (WM, yellow) used for analysis. **(G-H)** MTR was reduced in **(G)** GM and the **(H)** WM skeleton for P-Myrf(-/-) animals compared to control animals, regardless of wheel condition (FW, P-Myrf(+/-), n=25,, P-Myrf(-/-), n=25). Diffusion-weighted parameters, such as FA and MD, did not reveal any statistically significant differences. See Suppl. Fig. 3 for more details. Data presented as Mean±95%CI or boxplots. M1:Primary Motor Cortex, M2:Secondary Motor Cortex, S1: Primary Somatosensory Cortex, antCC: Anterior Corpus Callosum. Asterix indicate statistical significance (*p<0.05).

Brain microstructure was assessed *ex vivo* using a 9.4 T horizontal bore MR scanner (Varian, Palo Alto, CA, USA). Magnetisation transfer imaging and diffusion-weighted imaging were acquired. Three parameters were extracted (see Suppl. Fig. 2): i) magnetisation transfer ratio (MTR), which allows indirect detection of water bound to macromolecules, such as lipids and proteins, and is thus sensitive to myelin (Deloire-Grassin et al. 2000); ii) fractional anisotropy (FA), which describes the anisotropy of diffusion of water molecules; and iii) mean diffusivity (MD), which describes the rotationally invariant magnitude of water diffusion within brain tissue. FA and MD are sensitive but not specific to a number of white matter features, such as myelin, axon calibre, density and organisation, among others (Lazari and Lipp, 2021; Sampaio-Baptista and Johansen-Berg, 2017; Zatorre et al., 2012).

For a region of interest (ROI) analysis, five brain areas hypothesised to be involved in learning a running wheel task, or sensitive to exercise on the wheel, were selected: primary (M1) and secondary (M2) motor cortex, primary sensory cortex (S1), hippocampus (Hipp) and an anterior region of the corpus callosum (antCC), as illustrated in Fig. 3C. For exploratory purposes, we ran voxel-wise analyses to test for more local effects in any brain area across the grey matter mask and white matter skeleton (Fig. 3F). Finally, to investigate global changes, we tested for effects averaged across the whole grey matter (GM) and whole white matter (WM) skeleton (Fig. 3G,H).

#### Effects of genotype

First, to investigate the effect of the genetic manipulation on brain microstructure without considering behavioural experience of wheel running, we tested for effects of genotype in animals that were only exposed to the fixed wheel (FW, our control condition, n=25). When testing across the four grey matter regions of interest (ROIs) (M1, M2, S1, Hipp), P-Myrf^(-/-)^(n=13) were found to have significantly lower MTR, compared to P-Myrf^(+/-)^ (n=12) (mixed ANOVA; main effect of genotype (F(1,17)=7.32, *p*=0.015). Post hoc tests for individual ROIs (ANOVA, Bonferroni adjusted for number of ROIs) found a significant reduction in MTR in M1 (Fig. 3D, *p*=0.02) and M2 (Fig. 3D, *p*=0.032). Additionally, for the white matter region of the anterior corpus callosum (antCC), P-Myrf^(-/-)^ had significantly lower MTR, compared to P-Myrf^(+/-)^ (Fig. 3E, *p*=0.013). No statistically significant changes between groups were observed for diffusion-derived metrics of MD and FA (Suppl. Fig. 3). No effects were found in the voxel-wise analysis.

#### Effects of wheel condition

Next, to investigate the effect that running the complex wheel for 12 days may have on brain microstructure, we tested for effects of wheel condition (fixed vs complex wheel) in P-Myrf^(+/-)^ control animals only (CW:n=13, FW: n=12). No statistically significant difference between the two running wheel groups were detected in MTR, FA or MD in the ROI analysis (Suppl. Fig 3). A voxel-wise analysis revealed an increase in MD in the left M1 grey matter region (Suppl. Fig. 4A & B) for P-Myrf^(+/-)^ that ran the complex wheel.

#### Interactions between genotype and wheel condition

Finally, to investigate potential interactions between genetic manipulation and behavioural experience, we performed an ANOVA with genotype and wheel (CW & FW) as factors on all samples. Similar to effects found when considering the FW groups alone, significant main effects of genotype on MTR were found within specific grey matter and white matter ROIs (M1, M2, S1, antCC; Supp. Fig. 3A-E), as well as across the whole grey matter mask (Fig. 3G; F(1, 34)=7.77, *p*=0.019) and the white matter skeleton (Fig. 3H; F(1,34)=4.73, 0.037)., with P-Myrf^(-/-)^ animals having significantly lower MTR. However, no main effect of wheel or interactions between genotype and wheel on MTR were found. No main effects or interactions were detected in FA or MD in the ROI analysis (Suppl. Fig. 3F-N). The voxel wise analysis detected small clusters of changes in MTR between genotypes in cortical and midbrain regions (Suppl. Fig. 4C), yet no interaction between genotype and wheel was found for any voxel-wise analyses.

#### Correlations between MRI and behavioural performance

A voxel-wise analysis across all animals in the CW condition (n=26) tested for correlations between brain metrics and performance metrics (maximum speed & total distance run over 8 days). No relationship was found for MTR or FA. We found a negative correlation between distance run and MD in broad areas in cortical grey matter and basal ganglia, independent of genotype (Suppl. Fig. 3D&E). Mice that ran longer distances had lower MD, indicating that water in these areas is more restricted (regardless of direction), potentially indicating higher brain tissue density.

Taken together, these results indicate that the ablation of oligodendrogenesis in young adult mice for 5-6 weeks led to decreases in the myelin sensitive metric MTR, across grey matter and white matter regions. However, we found no interactions between wheel running and genotype, suggesting no evidence for changes in brain microstructure in response to wheel running that depend on oligodendrogenesis.

### Ablation of oligodendrogenesis changes oscillatory brain activity in Myrf-cKO mice in vivo

Previous work has indicated that even subtle deficits in myelination can lead to altered neuronal activity and synchronicity (Dubey et al., 2022; Gould et al., 2018; Kato et al., 2020), while modelling approaches indicate a role of activity-dependent myelination in oscillatory self-organization and homeostatic coordination of brain networks (Noori et al., 2020; Talidou et al., 2022).

As our histological and MRI data suggests altered myelination in P-Myrf^(-/-)^ mice, we tested oscillatory brain activity using cortical EEG recordings in freely-behaving P-Myrf mice, not exposed to running wheels. In each animal, the electrodes were implanted in two cortical regions (see Fig. 4A) at the age of P110-120, 5-6 weeks after tamoxifen treatment (Fig. 4B). Mice were individually housed in standard conditions. Data were recorded over a 2-day period, 1-2 weeks after surgery (P-Myrf^(+/-)^: n=3, P-Myrf^(-/-)^: n=4) during spontaneous sleep and waking in undisturbed animals.

**Fig. 4.**
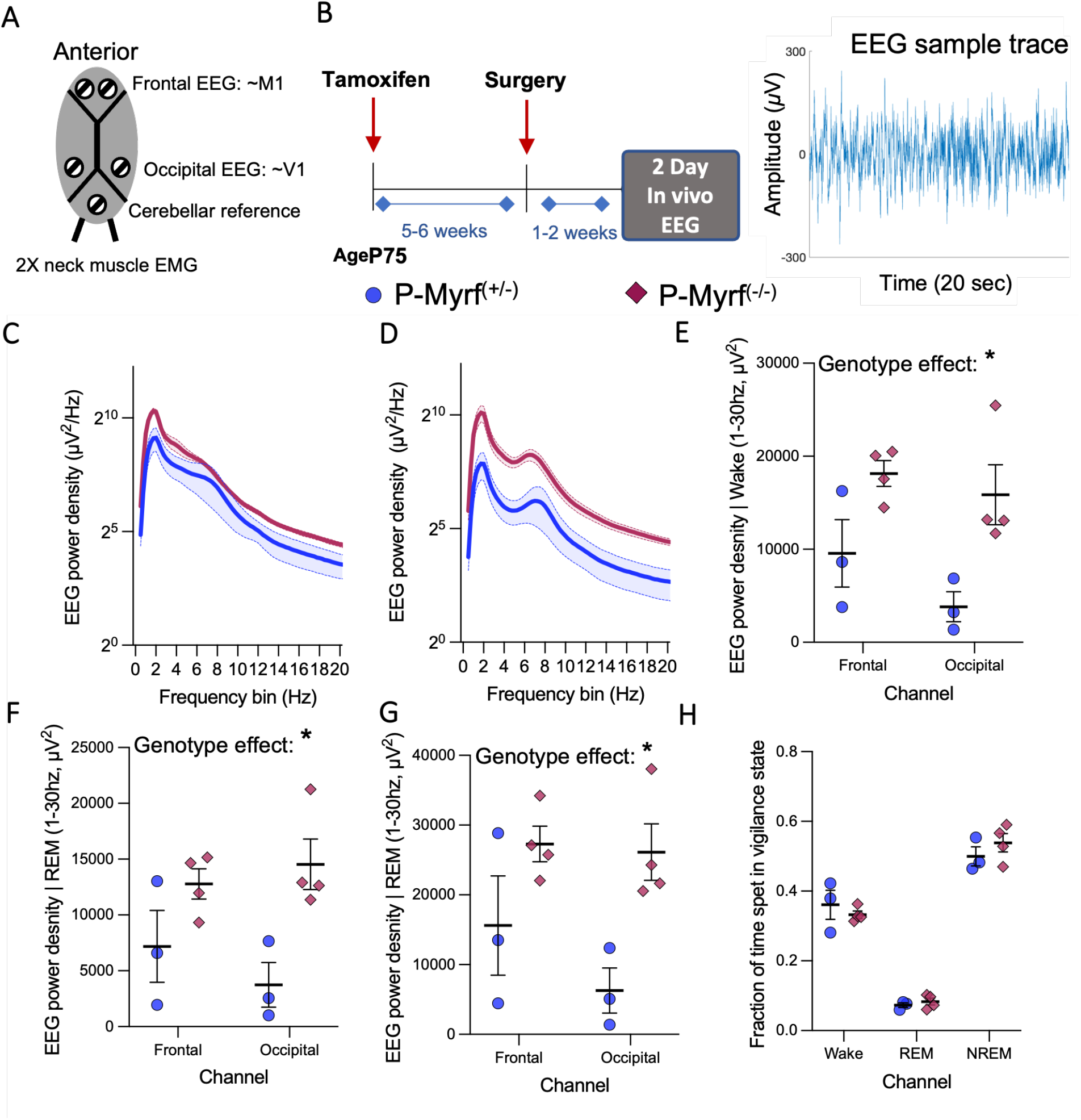
Ablation of oligodendrogenesis alters brain activity in the absence wheel running. **(A)** Schematic illustration of the location of screws implanted for in vivo EEG recording. **(B)** Schematic illustration of experimental design. **(C-D)** Mean group Power Spectral Density (PSD) by Frequency during wakefulness in **(C)** frontal channel and **(D)** occipital channels. Data presented as Mean±SED (**E**) The mean PSD across the 1-30 hz frequency bands was lower for animals with ablated oligodendrogenesis in the frontal and occipital channel during wakefulness. This genotype difference was found to be statistically significant across frontal and occipital channels (Mixed ANOVA) during wakefulness. **(F-G)** The effect was not specific to wakefulness, as statistically significant differences in mean PSD were also observed during (**F**) REM sleep and **(G)** non-REM sleep. **(H)** The fraction of time animals spent in different vigilance states. Recordings were undertaken in seven male adult mice (P-Myrf(-/-): n=4; P-Myrf(+/-): n=3). Data presented as Mean±SD. Asterix indicate statistical significance (*p<0.05).

Spectral analysis revealed a significant increase in EEG spectral power across broad frequency bands in P-Myrf^(-/-)^ animals in the frontal and occipital region during wakefulness (Fig. 4C-E), when compared to P-Myrf^(+/-)^ animals. Using a Mixed ANOVA (Main effect: genotype, within-subject factor: electrode location), the genotype difference in total EEG power density in 1-30 Hz frequency range was statistically significant across both cortical derivations during wakefulness (main effect for genotype: F(1,5)=10.4, *p*=0.023). This change in total spectral power induced by the conditional deletion of *Myrf in OPCs* was also observed during other vigilance states such as REM sleep (Fig. 4F, Suppl. Fig A&B; F(1,5)=12.7, *p*=0.016) and nonREM sleep (Fig. 4G, Suppl. Fig. C&D,F(1,5)=11.4, *p*=0.02). The fraction of time animals spent in different vigilance states did not differ between genotypes (Fig. 4H). The peak EEG power density during wake and REM sleep (0.25 hz resolution) in the Theta range (4-12 hz) was significantly different between genotype groups (F(1,5)=7.4, *p*=0.041, Suppl. Fig. 5E), yet the frequency of the peak was not (Suppl. Fig 5F). Altogether, these results provide evidence for changes in brain electrophysiology induced by the ablation of oligodendrogenesis for 7-8 weeks prior to EEG recording in freely moving and behaviourally naïve mice.

In summary, we replicated the previously reported behavioural phenotype of P-Myrf^(-/-)^ showing reduced capacity to improve performance over time on the complex wheel, which is prominent within the first hours of running. We additionally found a high degree of variation in engagement with the wheel, which should be taken into account when interpreting results. Further, our results suggest that ablation of oligodendrogenesis alters brain microstructure, as detected by a reduction in MTR, and brain activity, as indicated by increase in EEG mean spectral power, independently of behavioural experience on the complex running wheel. This suggests ablation of oligodendrogenesis in P-Myrf^(-/-)^ can itself lead to physiological and structural alterations in the brain.

## Discussion

This study investigated the consequences of disrupting OL differentiation during adulthood (Fig. 1) on behaviour (Fig. 2), brain structure (Fig. 3) and brain activity (Fig. 4). Our results replicated the key finding that ability to perform a complex wheel (CW) running task is disrupted by ablation of oligodendrogenesis (Fig. 2, McKenzie et al., 2014; Xiao et al., 2016). These group differences on the complex wheel were present within hours of running on the wheel. Additionally, using *ex vivo* MRI (Fig. 3) and *in vivo* EEG (Fig. 4), we found that the ablation of oligodendrogenesis altered brain microstructure and activity independently of the behavioural experience of running the complex wheel. Taken together, our findings indicate that inhibiting the formation of new oligodendrocytes in adult mice not only led to a decline in performance on the complex wheel but also brought about alterations in both the microstructure and activity of the brain, irrespective of animals’ experience with the task. Whether and how these task-independent changes in brain microstructure and function contributed to the observed behavioural phenotype requires further investigation.

### Replication of behavioural deficit on the complex wheel

We found large variability in the time animals spent engaging with the complex wheel task, which was in turn related to their overall performance (Fig. 2I). Accounting for this variability by analysing performance on the CW as a function of time animals spent on the CW (Fig. 2L-O) revealed that P-Myrf^(-/-)^ animal’s deficit on the wheel arises within the first hour of running (Fig. 2L-<).

However, it is difficult to clearly define the underlying deficit in P-Myrf^(-/-)^ mice that leads to the observed deficit on the CW, as the task demands of the CW are multifaceted. Running speed improvements over several days are observed on both the normal and complex wheel and can therefore reflect generic aspects of wheel exposure as well as specific demands of the complex wheel (Suppl. Fig 1D, Suppl. Fig. 6). Switching from a normal wheel to a complex wheel leads to a temporary reduction in running performance in WT mice (Suppl. Fig. 6, Hibbits et al., 2009), indicating there is some specific task demand of the irregular spaced rungs. Yet, changing the rung sequence does not affect performance, indicating that mice adapt a strategy to cope with irregular spaced rungs in general and do not learn a specific sequence (Suppl. Fig 6). Together, this indicates that mice need to find a new strategy (e.g. adapt their gait pattern, McKenzie et al., 2014) to cope with irregular rungs and be able to run and improve their performance on the CW.

A deficit within the first hour running the complex wheel suggests that a key task requirement impaired in P-Myrf^(-/-)^ animals is probed by early attempts to perform the CW task, rather than by learning processes evolving over several hours or days. Although the CW has been used as a test of subtle deficits in complex motor *execution* (Hibbits et al., 2009; Liebetanz et al., 2007; Schalomon and Wahlsten, 2002), the fact that P-Myrf^(-/-)^ mice already trained on the CW prior to tamoxifen treatment have no deficits when re-exposed to the CW (McKenzie et a l., 2014), together with the absence of any execution deficit on other motor tasks (Suppl. Fig. 1A-F, McKenzie et al., 2014), suggests that impairment that we and others have found in CW performance reflects impairment in the acquisition of a new skill rather than skill execution or general motor performance.

### Task-independent changes to brain structure and function might contribute behavioural deficit

How an inability to form new oligodendrocytes can cause the observed learning phenotype of P-Myrf^(-/-)^ mice within just a few hours is unclear. Previous studies suggest that rapid production of *Enpp6*-expressing immature oligodendrocytes can be observed within the first hours of exposure to the complex wheel, potentially contributing to circuit functioning and neuronal metabolism (Xiao et al., 2016). Yet, our understanding of how new oligodendrocytes contribute to skill learning and memory formation remains incomplete, with apparently different timescales of involvement found with different learning paradigms.

For example, while early learning deficits have been found with complex wheel running, studies investigating spatial learning and fear conditioning in mice with a conditional knockout of *Myrf* in OPCs found no deficit in early memory formation, but a deficit of memory consolidation over days or weeks (Pan et al., 2020; Steadman et al., 2020). Different aspects of myelin plasticity might occur over distinct timescales. For example, a study investigating motor learning in the dextrous reaching task found modulation of existing myelin in upper layers of M1 predominant in early stages of learning, while addition of new myelin sheaths was more relevant post learning (Bacmeister et al., 2022).

An alternative explanation for the behavioural deficit is that the disruption of oligodendrogenesis for weeks prior to behaviour testing in P-Myrf^(-/-)^ caused subtle changes in brain microstructure (Fig. 3) and activity (Fig. 4) which then impacted on capacity to learn a new skill. New oligodendrocytes (OLs) continue to differentiate throughout early adulthood (Hill et al., 2018; Hughes et al., 2018; Rivers et al., 2008; Young et al., 2013) and interfering with such ongoing cellular dynamics can have unintended consequences, as suggested by the current study (Fig. 3 & Fig. 4). Activity-regulated myelination, including oligodendrogenesis, is proposed to play a role in homeostatic coordination and oscillatory self-organization in local and large-scale brain networks (Dubey et al., 2022; Noori et al., 2020; Talidou et al., 2022) and mild impairments in myelination have been linked to decreased motor learning by increasing asynchrony and spontaneity in neural activity (Kato et al., 2020). Hence, it is possible that brain differences between genotype groups at the start of behavioural testing, as well as a lack of activity-dependent up-regulation of oligodendrogenesis triggered by exposure to the CW, might have combined to contribute to the observed behavioural phenotype.

### Changes in brain microstructure due to Myrf knock-out

When investigating the consequences of genetic ablation of OL differentiation on brain microstructure using *ex vivo* MRI, we found significantly lower MTR in multiple brain regions (Fig. 3). We hypothesised that disruption of oligodendrogenesis will affect brain microstructure, as it disrupts ongoing cellular dynamics and myelin formation. As MTR is related to myelin content of tissue (Lazari and Lipp, 2021), lower MTR may reflect reduction in myelination caused by the significant reduction in new oligodendrocyte formation (Fig. 1) induced for ∼5-6 weeks prior to *ex vivo* MRI. However, estimating how much myelin content was added during the timeframe of interest is difficult and requires further histological investigation. Metrics derived from diffusion-weighted imaging (FA, MD), that are sensitive to tissue microstructure and less sensitive or specific to myelin, were not significantly affected by the deletion of *Myrf* (Suppl. Fig. 3). The finding that changes in MTR were widespread across WM and GM regions is consistent with the global ablation of OL differentiation across the whole CNS.

We found no evidence for task-dependent changes in brain microstructure related to myelination when investigating how motor learning on the complex wheel affected brain microstructure (in comparison to a fixed wheel control condition). Specifically, we found no effect of complex wheel running on the myelin-sensitive metric MTR (Fig. 3, Suppl. Fig. 3), regardless of whether animals had disrupted oligodendrogenesis. We had hypothesised that motor learning should lead to changes in myelin sensitive metrics of brain microstructure if associated with increase in myelination, as previously reported in the context of the skilled reaching motor learning tasks in rats (Sampaio-Baptista et al., 2013). However, there are several important methodological differences between these studies, including species (rat vs mouse) and training task. The reaching task involves the development of fine motor movements in comparison to the gross motor movements necessary to perform the CW task. Furthermore, MRI techniques employed here can only measure cellular effects very indirectly and are less sensitive to small effects compared to histological approaches employed by others, which showed increased oligodendrocytes differentiation in motor cortex and subcortical WM due to CW running (McKenzie et al., 2014; Xiao et al., 2016). In addition, the fixed wheel control condition we employed might have offered more environmental enrichment in comparison to home cage controls that are typically used. Furthermore, it is possible that at the timepoint of our measurement, immediately after 12 days of training on the complex wheel, experience-induced changes in myelination might not have been prominent enough to cause detectable changes in MTR. Together, we did not find evidence for learning-induced changes in myelin sensitive MRI measures of brain microstructure after learning to run the complex wheel.

### Electrophysiological changes cause by Myrf knockout

In addition to genotype differences in microstructure, deletion of *Myrf* in OPCs caused alterations in oscillatory brain activity (Fig. 4), as reflected in higher EEG power density across broad frequency bands. Increases in spectral power of LFPs have been reported in Plp1-KO models with mild myelin deficits (Gould et al., 2018) and cuprizone models of demyelination (Dubey et al., 2022). Associated computational modelling indicated that a decrease in conduction velocity, as a result of reduced myelination, can lead to an increase in spectral power (Gould et al., 2018). Additionally, adaptive myelination, disrupted in P-Myrf^(-/-)^ animals, might regulate homeostatic coordination and oscillatory self-organization in local and large-scale brain networks (Dubey et al., 2022; Noori et al., 2020; Talidou et al., 2022). Hence, it is plausible that reduced myelination and disruption of adaptive myelination, caused by ablated oligodendrogenesis over a 6-7 week period, led to the observed increases in EEG power density. However, identification of the underlying mechanism driving such broad alterations in brain activity requires further investigation.

### Summary and conclusion

In summary, the findings presented in this study suggest that ablation of oligodendrogenesis by conditional deletion of Myrf in PDGFa-positive OPCs leads to subtle but detectable physiological and structural differences in the brain. The previously detected, and here replicated, behavioural deficit on the complex running wheel might therefore partly arise as a result of these task-independent brain differences, in addition to impairments in oligodendrogenesis driven by learning.

The current findings raise many questions for future work. For example, the timescale over which these structural and physiological changes develop should be explored as we only sampled effects at a single timepoint. Further, it remains to be tested whether the effects found here hold for other transgenic models of disrupted oligodendrogenesis. While the use of such models has thus far been predominantly to study of effects on learning, future work could investigate whether and how continuous oligodendrogenesis plays a broader role in maintaining brain microstructure and function.

This study suggests that the implications of disrupting oligodendrogenesis in the adult mouse brain extend beyond specific task-related effects and should be taken into account when interpreting behavioural phenotypes and designing future experiments.

## Acknowledgements

The authors thank Eveliina Hanski and Luke Baxter for feedback on previous version of the manuscript. The authors also want to thank Piergiorgo Salva and the entire Plasticity Group at the Wellcome Centre for Integrative Neuroimaging for constructive discussions and suggestions during the project.

## Materials and methods

### Animals

All work carried out conformed to UK Home Office legislation (Scientific Procedures Act 1986).

### Mouse breeding and genetic background

Mice used in this study were originally generated in the Richardson laboratory (McKenzie et al., 2014), UCL by crossing *Pdgfra-CreER^T2^* transgenic mice (Rivers et al., 2008) with Cre-conditional reporter mice *Rosa26R-YFP* (*R26R-YFP*) (Srinivas et al., 2001) to generate double-homozygous offspring. *Cre* and *YFP* reporter lines in the Richardson Laboratory were maintained separately by crossing (more than 10 generations) with C57B6 females (Charles River; Margate, UK). These *Pdgfra-CreERT2 :Rosa26R-YFP* were crossed with *Myrf ^(fl/fl)^* mice (Emery et al., 2009) to obtain *Pdgfra-CreERT2: R26R: Myrf* ^(+/fl)^ offspring, which were sibling-mated to obtain *T2 Myrf ^(fl/fl)^* and *Myrf* ^(+/fl)^ littermates. These animals were finally crossed to generate cohorts of *Pdgfra-CreER : (fl/fl) (+/fl) R26R-YFP: Myrf* and *Myrf* littermates for behavioural testing and further breeding. In the Richardson Laboratory, *Myrf ^(fl/fl)^* mice were obtained originally (in 2010) on a mixed 129/CBA/C57B6 background. *T2 T2* Five breeding pairs of *Pdgfra-CreER^T2^ : R26R-YFP: Myrf ^(fl/fl)^* and *Pdgfra-CreER^T2^ : R26R-YFP: Myrf* ^(+/fl)^ (referred to as P-Myrf ^(*fl*/*fl*)^and P-Myrf ^(+/*fl*)^) were sent to our laboratory from the Richardson Lab, UCL in 2016 and were used to breed animals for our study by crossing P-Myrf ^(*fl*/*fl*)^ with P-Myrf ^(+/*fl*)^ mates. Mice carried copies of *Pdgfra-CreER^T2^* on both alleles and were either *R26R-YFP ^(+/-)^* or *R26R-YFP ^(+/+)^*. Two generations of animals were bred before behaviour testing started for the current studies and animals from a total of 4 consecutive generations were used for behaviour testing. Hence, P-Myrf^(*fl*/*fl*)^ and P-Myrf ^(+/*fl*)^ were maintained by sibling crossing for 6 generations in our facilities.

For testing the normal and complex wheel running in wild-type mice, twenty C57BL/6JOlaHsd mice (8 males and 8 females) were purchased from Envigo, UK at age P60. The mice were given a one-week period to acclimate before commencing behavioural testing.

### Mouse Genotyping

Genotyping for Cre-ER^T2^ (Cre 5’-TCG ATG CAA CGA GTG ATG AG and Cre 3’-TTC GGC TAT ACG TAA CAG GG; Product Size 481bp) and R26R-YFP (R26WTF1-AAA GTC GCT CTG AGT TGT TAT, R26WTR1-GGA GCG GGA GAA ATG GAT ATG, R26KOF1-GCG AAG AGT TTG TCC TCA ACC) was done by PCR amplification of genomic DNA. Myrf ^(*fl*/*fl*)^ mice were genotyped by PCR amplification of genomic DNA using primers within intron 7 flanking the first lox site (5’-AGGAGTGTTGTGGGAA-GTGG and 5’-CCCAGGCTGAAGATGGAATA), which gives a 281 bp product for the wild-type allele and a 489 bp product for the *loxP*-flanked allele (Emery et al., 2009).

### Tamoxifen administration

Tamoxifen (Sigma T5648) was dissolved at 40 mg/ml in corn oil on a shaker overnight at RT and by subsequent sonicating at 21°C for one hour. P-Myrf ^(+/-)^ and P-Myrf ^(-/-)^ mice were generated by administering tamoxifen (0.3 mg/g body weight in corn oil) by oral gavage for four consecutive days to *Pdgfra-T2 (flox/flox) T2 (+/flox) CreER :R26R-YFP:Myrf*and *Pdgfra-CreER :R26R-YFP:Myrf*. Animals were left for at least 2 weeks before experimental procedures. During, and for a week after tamoxifen treatment, nutritious jelly and mash were provided for all mice to prevent excessive weight loss due to the adverse effects of the drug.

### Behaviour testing of mice

Assessment of behavioural phenotype was conducted by the same researcher. Wherever possible, equal numbers of animals of each sex were included in each experimental group. For experiments for which data collection was not automated, researchers were blinded to the genotype of the animals during data collection and analysis. All mice were maintained on a 12-hour artificial light-dark cycle (9:00-21:00). Food and water were provided *ad libitum*.

### Complex running wheel

Cages with running wheels were purchased from Lafayette Neuroscience (Scurry mouse mis-step wheel Model 80821S). This voluntary running wheel cage comes equipped with a wheel (circumference = 0.389 m) that allows individual rungs to be removed to create a complex wheel or rung configuration (Liebetanz et al., 2007; McKenzie et al., 2014). The mice were provided with a “complex wheel” (CW) with 16 out of 38 rungs removed to create a wheel with an irregular pattern of 22-rungs (Fig. 2A), a configuration that was adapted from (McKenzie et al., 2014). For behaviour testing, mice were single caged with the running wheel, a paper house filled with nesting material and wood chippings to cover the floor of the cage. Mice were accustomed to the cage for 30 mins before being provided access to the running wheel by removing a dividing wall to the compartment where the wheel was located. Mice were placed in the wheel cages up to 2 hours before the start of the dark period. Mice ran spontaneously, without artificial reward. Food and water were available *ad libitum* within the specialised running wheel cages.

Infrared sensors recorded wheel rotation, which allowed computer-controlled recording of wheel rotation as a function of time for 20 cages simultaneously for 24h a day (The Scurry Activity Monitoring Software, Lafayette Neuroscience, Model 86165). For data analysis, counts of full wheel rotations were exported into a spreadsheet, divided into time bins of 10 seconds. Custom R scripts were used extract the following running parameters from the baseline activity data: 1) distance travelled, which represents the total number of wheel rotations × wheel circumference. 2) Active intervals, which represent the number of 10 second intervals in which at least one full wheel rotation was recorded. 3) Average speed, which represents the sum of the distance run divided by the number of active intervals. 4) Maximum speed, which was calculated by determining the average speed in the 1% of fastest intervals run. Analysis of performance was conducted for either chronological time intervals (length of time vector) or for intervals of time that animals spent running on the wheel (length of vector containing only active intervals on the wheel).

Running speed of animals (distance/time) was conducted using time bins of 10 seconds. McKenzie *et al*. (2014) calculated the average running speed by quantifying wheel spins during a 1 min time bin. Yet, in the current study between 35-40% of continuous running bouts were found to be shorter than 1 minute long (data not shown). That meant that average running speed was often not accurately estimated when using 1 min intervals to calculate speed. Hence, 10 sec bins were chosen for this study.

### Complex running wheel and fixed wheel testing for ex vivo MRI experiment

A total of 24 animals were tested on the complex running wheel (P-Myrf^(+/-)^:n=12, 5 females; P-Myrf^(-/-)^:n=12, 4 females) in specific running wheel cages as described above. Additionally, 25 littermates were exposed to a static fixed complex wheel that was not able to rotate (referred to as fixed wheel, FW) in cages with otherwise identical set up (P-Myrf^(+/-)^:n=12, 4 females; P-Myrf^(-/-)^:n=13, 5 females). As the fixed wheel had the same missing rungs pattern that allows animals to pass between rungs, animals could climb and interact with the wheel as a kind of environmental enrichment yet could not run and turn the wheel. The experiment consisted of 5 distinct batches of P-Myrf animals bred from the same set of 5 breeding pairs over consecutive litters. Littermates were semi-randomly assigned to cages equipped with either a complex wheel (CW) or a fixed wheel (FW) and were tested in the same environment and timeframe. Semi-random assignment meant that animals were randomly assigned to a wheel type, while attempting to balance the numbers of genotype, sex and littermates equally between experimental groups.

### Accelerating RotaRod

Mice were tested on a standard RotaRod apparatus (Ugo Basile, Italy). Mice were tested for their ability to stay on a rod 3 cm in dimeter (Suppl. Fig. 1 B). The apparatus allowed simultaneous testing of up to 5 mice, yet a maximum of 3 mice were tested at the same time. Mice were habituated to the testing room for 30 mins before behaviour testing, by placing their cages into the room. Mice were trained for three trials on the rotarod at a constant speed (4 rpm), 120 s each trial, 2 days before behaviour testing, at which speed all mice stayed on the rod for the duration of the trial.

On the day of testing, each mouse was held by the tail and allowed to grasp the rotating rod. If it was still in place after 10 s, then the rotation of the rod was accelerated by 20 rpm to a maximum speed of 40 rpm. The time at which each mouse fell off the rod was recorded, starting from the moment of acceleration. If a mouse managed to stay on the rod for longer than 150 s, the trial was stopped, and a time of 150 s was recorded. Each mouse was tested 3 times on the same day. Animals were allowed to rest for 10 minutes between trials.

### Balance beam

The beam apparatus consists of 1 meter round smooth wooden beams of 15 mm or 7.5 mm diameter positioned 50 cm above the table. A soft foam matt was placed beneath the beam to cushion any potential fall. The beam rested on the edge of a wooden box on one side, at which animals were placed for testing. On the other side, the beam rested on the edge of the animal’s home cage (Suppl. Fig. 1C). The beam was slightly angled (∼10 degrees) so that animals had to walk uphill to reach their home cage. A lamp (with a 60-watt light bulb) was used to shine light above the start point and serve as an aversive stimulus, and the light inside the room was slightly dimmed. Mice were habituated to the testing room for 1 hour before behaviour testing, by placing their cage inside the room. Three days before behaviour assessment on the balance beam, mice were trained to cross the beam. Training was done by placing the mice on the beam (1.5 cm diameter) opposite the home cage. During training mice were freely allowed to cross the beam and reach their home cage. This procedure was repeated 3 times with a 1-min rest time between attempts. Training was conducted as almost all mice stopped on the beam or tried to turn around in the middle during the first exposure to the apparatus. Only after learning the location of their home cage during training do the mice reliably cross the beam without stopping and turning around, which allows reliable and comparable data collection. All mice successfully crossed the beams during training at least one time.

For testing, mice were placed on the beam and freely allowed to cross the beam. Mice were first tested on the wider beam (1.5 cm diameter). After a 5-min resting time in the home cage, the beam was exchanged for the narrow (0.75 cm diameter) beam and the same animal was tested again. All animals successfully crossed the beams without falling. The apparatus was cleaned with ethanol (70%) before testing another animal.

Testing was recorded using an HD camera placed on the side of the beam opposite the home cage. Videos were analysed by a blinded experimenter. Foot slips (defined as the foot coming off the top of the beam) and the number of steps taken to cross the beam were quantified.

### Perfusion of animals and tissue processing

Animals were deeply anaesthetized (Euthatal: 80mg/kg, i.p) with sodium pentobarbital and transcardially perfused with 10 ml PBS, followed by 30 ml paraformaldehyde (4% w/v in 0.1 M phosphate buffer) with the help of a perfusion pump (Flow rate: 4ml/min). After perfusion, animal skulls were dissected by removing soft tissue from the outside of the skull (skin, muscle, eyes, etc.) and the lower jaws. Dissected skulls containing the CNS tissue were stored overnight in 4% PFA at 4°C for 24h. Samples were then washed with phosphate buffer saline (PBS) before being stored in PBS at 4°C. Samples were sent to the Center for Image Sciences, UMCU, Utrecht, Netherlands for magnetic resonance imaging (MRI). After MR imaging, tissue samples were sent back to the University of Oxford, cryoprotected with 30% sucrose solution, embedded in OCT and stored at -80°C until processed for histological analysis.

### Immunohistochemistry of free-floating brain sections

For Immunohistochemical analysis, coronal cryosections (30 μm) were cut (anterior to posterior) from the first section at which the anterior corpus callosum was visible and continuously connected from one hemisphere to the other. Brain sections were collected and washed in Tris-buffered Saline (TBS; Sigma) and processed as floating sections. All incubations and washes were performed on a shaker at 60 rpm. Sections were blocked in TBS containing 10% Normal Goat Serum (Sigma), 0.5% Triton-X (Sigma) and 1% BSA for 2 hours at room temperature. The primary antibodies used were anti-PDGF-Ra (rabbit, New England Biolabs, 1:400 dilution), anti-APC (clone CC1; mouse, Calbiochem, 1:200 dilution) and/or anti-GFP(YFP) (chicken, Avas, 1:500), which were diluted in TBS containing 10% of the blocking solution (see above) at 4°C. Samples were then incubated with corresponding fluorophore conjugated secondary antibodies (1:500, Alexa FluorO-488 or 546, Thermo Fischer Scientific), diluted in blocking solution (1:10) for 2h at room temperature. Samples were mounted and dried for ∼10 mins before being sealed under a glass cover with Vectorshield anti-fade mounting medium (Vectorshield laboratories) and viewed by a Zeiss L subse SM-700 confocal microscope and ZEN software. Cells were counted in low-magnification photo-micrographs (x20 objective) of non-overlapping fields of coronal sections (320mmx320mm each) of the corpus callosum, between the dorsolateral corners of the lateral ventricles (6 fields per section, three sections per animal), as illustrated in Suppl. Fig. 6. The surface area of corpus callosum was calculated for each animal to determine cellular density. Image analysis was performed using Fuji/ImageJ (NIH).

### Statistical analysis of histological and behavioural data

#### Criteria for excluding animals from complex wheel data analysis

Previous experiments reveal that a small subset of healthy untreated animals show unusually low engagement in voluntary wheel running. To prevent such outliners from skewing overall group performance, we set the following pre-determined exclusion criteria for analysis of complex wheel behaviour performance: animals were excluded from the analysis if: 1) they ran less than 1km in 8 Days (P-Myrf^(+/-)^; n=4, females; P-Myrf^(-/-)^; n=1), or 2) ran less than 100 meters per day for at least half of the days recorded (P-Myrf^(+/-)^; n=2, females; P-Myrf^(-/-)^; n=1). In total, that led to the exclusion of 8 animals based on these criteria (P-Myrf^(+/-)^; n=6; P-Myrf^(-/-)^; n=2).

#### Testing for the effect of covariates on complex wheel running

Before analysing the effect of genotype on behaviour, we assessed whether a number of different covariates might influence our findings, in order to determine which covariates to include in subsequent analyses. First, to test for a potential effect of the generations and sex on the performance on the CW, a mixed two-way ANOVA (within subject factor: days (over 8 days of running); between subject factors: sex, generation) was used to analyse daily running speed (average or maximum). This analysis revealed that sex, but not generation, had a significant effect maximum running speeds (sex: F(1,116)=22.68, p<0.001) on the complex wheel, with females consistently outperforming males (Suppl. Fig. 1G). Apart from consistently running longer distances on the CW (total distance in m: female, M±SD=35505±22374; males, M±SD=16413±14426), females also spent more time on the CW than males over a period of 8 consecutive days (Suppl. Fig. 1H). While there was an expected difference in body weight between sexes, there was no difference in body weight between P-Myrf^(−/−)^ and P-Myrf^(+/−)^ animals for either sex at the start of behaviour testing (body weight in g: male-Myrf^(+/)^, M±SD=24.1±3.4; male-Myrf^(-/-)^, M±SD=24.3±2.4; female-Myrf^(+/-)^, M±SD=19.7.1±1.4, female-Myrf^(-/-)^, M±SD=19.4±2.3).

The main analysis of behavioural performance on the CW combines data from several experiments. Hence, the age of tamoxifen treatment (range: P58-93, M±SD=77.11±8.6), the age at behaviour testing (range: P92-135, M±SD=109.9±7.9) and the days between tamoxifen treatment and behaviour testing (range: P21-43, M±SD=32.8±7.5) varied between animals, experiments and generations (Suppl. Table 1). However, we found no statistically significant correlation between the performance of the P-Myrf^(−/−)^ on the CW (average running speed) and any of the age parameters mentioned above in the compiled data set of all P-Myrf animals. Yet, it is of note that our experiments were not specifically designed to compare the effect of these age parameters on running performance.

In summary, of the potential covariates tested, only sex was found to be significantly associated with running behaviour and so this factor was included as a fixed factor in all subsequent analyses of genotype differences to control for the induced variability.

#### Statistical analysis method

Statistical analysis was performed using R (R Core Team, 2021), using he rstatix (0.7.0; Kassambara, 2021) and the psycho (v0.6.2; Makowski, 2021) packages. The statistical tests used for a particular analysis are mentioned during the data presentation in the result sections. Normality of the data and assumptions of homogeneity of variance were tested (e.g. with Levene’s Test) before using parametric statistical analysis. If these assumptions were not met, non-parametric alternative tests were used for statistical analysis, or the test parameters were adjusted to account for the lack of homogeneity of variances (e.g. using un-pooled variances and a correction to the degrees of freedom for unpaired t-test) whenever possible.

Performance on the complex wheel was analysed using a mixed two-way ANOVAs. However, sample data did not always meet the assumption of normality due to a tendency towards bimodal distribution in the sample population. Despite this we proceeded to apply ANOVA as I) this statistical analysis shows a certain robustness to type 1 error due to moderate violation of assumptions of normality at the range of sample sizes used in this study (Arnau et al., 2013; Blanca et al., 2018, 2017), II) no ideal alternative non-parametric statistical analysis exist for a mixed model ANOVA, and III) the same statistical analysis was used for the data set that we are attempting to replicate. If the assumption of sphericity (Mauchly’s Test) was violated in the mixed two-way ANOVA, Greenhouse-Geisser procedures were used to estimate epsilon in order to correct the degrees of freedom of the F-distribution. In this case, adjusted degrees of freedom are presented. Asterisks are used to indicate statistical significance when appropriate: *=p<0.05, **=p<0.01 ***=p<0.001. Details of statistical analysis approaches and results for each dataset are provided in Suppl. Table 2

### MRI methods

#### MRI acquisition

Post-mortem high-resolution structural MRI was performed on a 9.4 T horizontal bore MR system (Varian, Palo Alto, CA, USA) at the Center for Image Sciences, UMCU, Utrecht, Netherlands, equipped with a 6 cm ID gradient insert with gradients up to 1 T/m. A custom-made solenoid coil with an internal diameter of 2.6 cm was used for excitation and reception of the MR signal. Three perfusion-fixed brains were inserted with the skulls intact in a custom-made holder and immersed in non-magnetic oil (Fomblin, Solvay Solexis).

Diffusion weighted imaging (DWI) was acquired with 3D eight shot spin-echo EPI sequence with the following parameters: 60 diffusion encoding directions; 1 average per image; 5 images with no diffusion weighting; TR/TE 500/38.6 ms, field of view: 24×20×20 mm; isotropic resolution of 0.125 mm ; δ/Δ 5.5/8.91 ms, diffusion gradient strength (half-sine shape) 73 G/cm, b = 3522 s/mm2; total acquisition time 12.3 hrs.

Magnetization transfer images were also acquired with a 3D eight shot spin-echo EPI sequence with a train of saturating pulses in front of the image acquisition saturating frequencies 5 kHz, 10 kHz, and 100 kHz off resonance. Four volumes per frequency offset were acquired (TR/TE 500/28ms; field of view: 24×20×20 mm; resolution of 0.125 mm isotropic; total time 2.3 hrs).

A total of 71 animals were scanned for *ex vivo* MRI. Unfortunately, a subset of 20 animals were perfused by a different experimenter using a different perfusion method. Due to complications with the perfusion method and a significant deviation in measured MRI metrics from the remaining cohort, these 20 samples had to be excluded from the analysis.

#### MRI data processing

All data were Fourier transformed and combined using home-written software in MatLab (Mathworks®) from the Dijkhuizen lab, UMCU, Utrecht, Netherlands. These scripts can be provided on request. DWI data were analysed with the FMRIB Diffusion Toolbox. Voxel-wise values of fractional anisotropy (FA) and mean diffusivity (MD) were estimated from the original DWI data using ‘dtifit’.

For alignment of the images to the same space, individual brain images were digitally separated from the triplets in which they were acquired. For the optimal registration of images in grey and white matter, two different contrasts were combined (both weighted equally): i) FA and ii) and mean of all diffusion weighted directions. A study specific template of all subject images was then acquired using an automated image registration pipeline as described previously (Lerch et al., 2011, 2008). In short, this approach brings all scans into anatomical alignment in an automated and unbiased fashion using a combination of mni_autoreg tools (Collins et al., 1994) and Advanced Normalization Tools (ANTS) (Avants et al., 2008). FSL (for FMRIB Software Library) tools, version 6.0 (www.fmrib.ox.ac.uk/fsl) were used for linear (FLIRT) and nonlinear transformations (FNIRT) to the study specific template (Scripts can be provided on request). Transformations gained from this approach were applied to the individual modalities of interest (FA, MD, MTR).

White matter structures were analysed using a modified version of Tract-Based Spatial Statistics (TBSS; Smith et al., 2006). The skeleton for TBSS was thresholded at an FA value of 0.28 as to reliably contain major tracts in the mouse brain that can be accurately aligned across individuals. Finally, the FA values of the tract centers (i.e., maximum FA values) were projected onto the skeleton for each mouse brain and fed into statistical analysis. MD and MTR values for the same voxels were also projected into the skeleton.

#### MRI data analysis

For region of interest **(ROI)** analysis, an aggregate atlas combining 182 individually segmented structures (Dorr et al., 2008; Richards et al., 2011; Steadman et al., 2014; Ullmann et al., 2013) was registered to the unbiased consensus average of the current study (the atlas is available at http://repo.mouseimag-ing.ca/repo/DSURQE_40micron_nifti/) and masks for the specific ROI (M1, M2, S1, Hippocampus) were ex-tracted from the atlas. Mean values from each ROI for each subject were extracted for statistical analysis in R studio. Mixed ANOVAs were used for group comparison. where regions of Interest (ROIs) were within subject factors and genotype and/or wheel were between subject factors. As data was collected from 4 consecutive experimental groups, experimental group (n=4) was included as a fixed factor to control for introduced variability. Significant effects in within-subject factor (ROIs) were followed by a post-hoc simple comparison between groups (wheel or genotype), Bonferroni adjusted for number of ROIs. Statistical analysis was performed using R (R Core Team, 2021), using he rstatix (0.7.0; Kassambara, 2021). To check for outliers, mean values for each metric were extracted from the whole brain mask and checked for extreme outliers (‘*identify_outliers’*, *rstatix* package, R). One animal (P-Myrf^(-/-)^, CW) was identified as an extreme outlier and subsequently removed from the ROI analysis. Statistical analysis including the outlier are provided in Suppl Table 2.

For voxel-wise analysis across the whole brain, nonparametric permutation testing with a cluster-forming threshold of t > 2 and 5000 permutations were used to determine corrected p values. Clusters with a corrected significance of p < 0.05 were deemed significant. We tested for differences between the genotype groups (Myrf^(-/-)^ vs Myrf^(+/-)^, wheel type (complex wheel vs fixed wheel) and for a genotype*wheel type interaction (Scripts and GLMs can be provided on request). We also tested for voxel-wise for correlations between contrast and wheel performance (maximum speed, total distance).

Finally, to test for global effects, we calculate mean values for each MR metric across all voxels within the GM mask or the WM skeleton.

### EEG methods

#### Electrophysiology data collection

Chronic electrophysiological recordings were undertaken in seven male adult mice (P-Myrf ^(-/-)^: n=4; P-Myrf (+/-): n=3). All mice received tamoxifen at age P75 and were implanted at ages P110-120, and the reported recordings were collected at age P130. The animals were surgically implanted with a custom-made headstage tethered to electrodes for the continuous recording of electroencephalogram (EEG) and electromyogram (EMG). EEG screw electrodes (Fine Science Tools) were inserted into the skull. 5 electrodes were implanted in total: above the right and left frontal cortex (primary motor area: anteroposterior +2 mm, mediolateral +2 mm), above the right and left occipital cortex (primary visual area: anteroposterior +3.5 mm, mediolateral +2.5 mm), and above the cerebellum (which served as a reference). As per <u>Fisher *et al*., 2016)</u>, two custom-soldered stainless steel wires were inserted into the right and left nuchal muscles respectively, for the recording of EMG. A schematic diagram of implantation locations is shown in Fig. 4A.

Immediately before surgery, analgesics were administered (subcutaneous injection of 1–2 mg/kg metacam and 0.08 mg/kg vetergesic). Isoflurane was used for induction and maintenance of anesthesia throughout the surgical procedure (4% and 1–2% respectively). After surgery, analgesics were given for at least 3 days (1–2 mg/kg oral metacam) and animal wellbeing was closely monitored for 1-2 weeks, until the animal returned to baseline conditions. All procedures were performed under a UK Home Office Project License and conformed to the Animal Scientific Procedures Act 1986.

Two animals were implanted each day, with all surgeries taking place in the same week. Order of surgery was randomised to ensure balanced allocation of genotype to time of day of surgery (i.e. morning surgery vs afternoon surgery) and to time of week surgery (i.e. early in the week or late in the week), to avoid systemic bias in implantations and in length of time elapsed between implantation and recording.

#### Electrophysiology data processing

Data acquisition was performed using the Multichannel Neurophysiology Recording System (TDT, Alachua FL, USA). EEG/EMG data were filtered between 0.1–100 Hz, amplified (PZ5 NeuroDigitizer pre-amplifier, TDT Alachua FL, USA) and stored on a local computer at a sampling rate of 256.9 Hz. EEG/EMG data were resampled offline at a sampling rate of 256 Hz. Signal conversion was performed using custom-written Matlab (The MathWorks Inc, Natick, Massachusetts, USA) scripts and was then transformed into European Data Format (Fisher et al., 2016). Custom MATLAB scripts were used to estimate the power spectral density using the multitaper method (“pmtm()” function, Signal Processing Toolbox, Matlab) for 4-s epochs f. A 0.25 Hz resolution was used for plotting of EEG power density across frequency bands. Total power within the 1-30hz range across the frontal and occipital channel was calculated and compared using ANOVAs for repeated measures (Within-Subject Factor: Location; Between Subject Factor: Genotype). Peak frequency was defined as the frequency bin (0.25 Hz resolution) with the highest power. Power within specific frequency bands was calculated and compared between genotype groups: delta, δ (0.5–3 Hz), theta, θ (4– 12 Hz), beta, β (12.5–25 Hz), and gamma, γ (30–80 Hz). Statistical analysis was performed using R (R Core Team, 2021), using he rstatix (0.7.0; Kassambara, 2021).

#### Scoring and analysis of vigilance states

Recordings were quality-checked during acquisition. All data was manually scored by a blinded investigator (S.R.) for vigilance states, as previously described in Fisher et al. (2016). Briefly, waking was defined based on a low-amplitude fast-frequency EEG activity and high EMG amplitude, NREM sleep was characterised by the predominance of slow waves and sleep spindles on the EEG and low EMG tone, while REM sleep was characterised by strong theta EEG activity, especially in the occipital derivation and low EMG activity. To validate scoring accuracy, a subset of the data was re-scored by another investigator (A.L.). Scoring quality was further verified by a third investigator (V.V.). The channel of the left hemisphere was selected for group comparison, unless a blinded investigator (V.V.) indicated that the quality of the data was significantly better in the right hemisphere, in which case the right hemisphere channel was used.

## Supplementary Material

**Suppl. Fig. 1.**
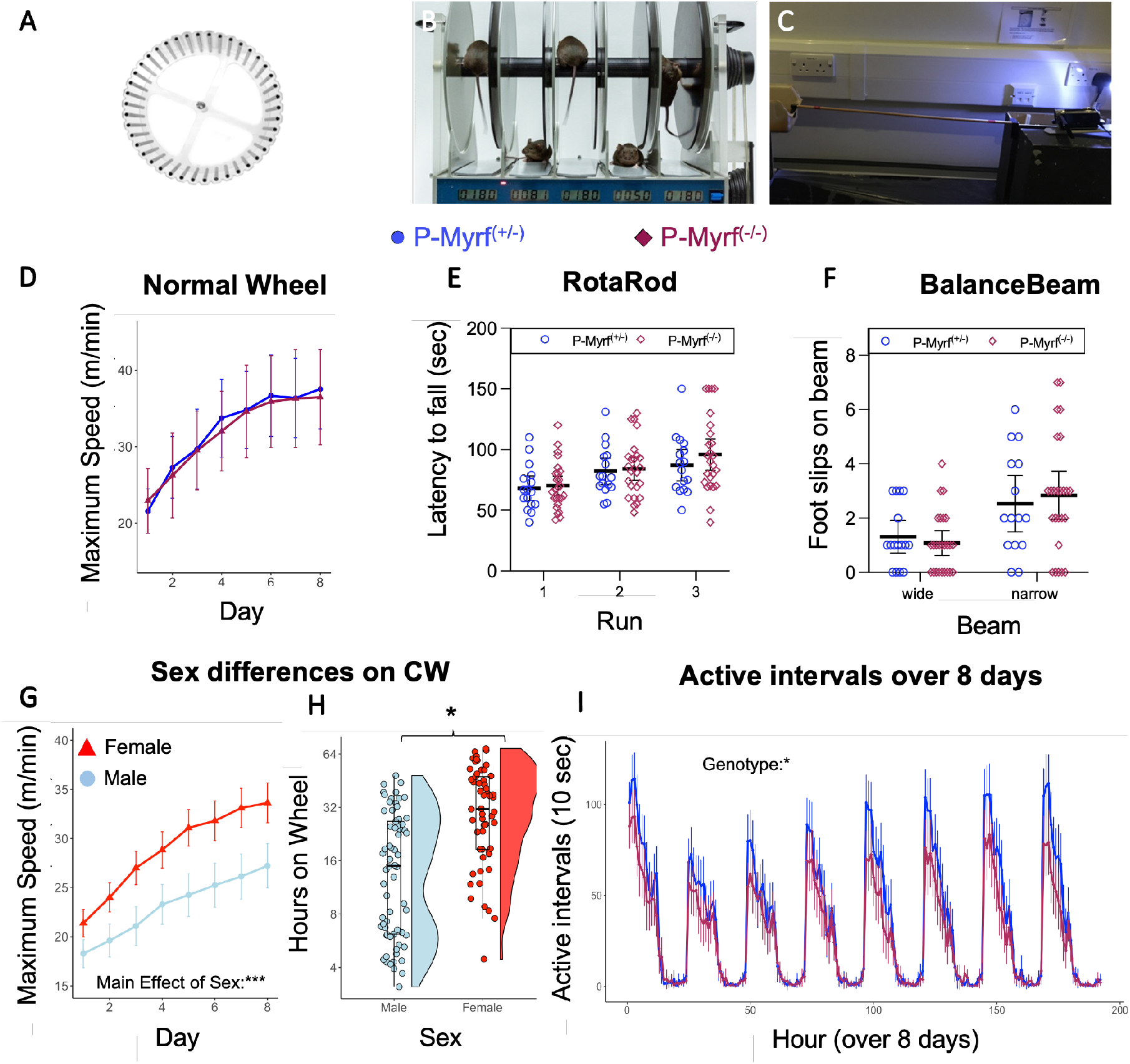
Control task for motor behaviour and supplementary data for the complex running wheel data. **(A)** To test general performance on a running wheel, a cohort of P-Myrf(-/-) and P-Myrf(+/-) animals were tested on a normal wheel which had none of the rungs removed. To test for general motor ability and coordination, a cohort of animals was tested on the (B) RotaRod and (C) Balance Beam task. (D) Conditional knockout of Myrf did not affect performance on a normal running wheel (P-Myrf(+/-); n=17; P-Myrf(-/-); n=18), nor did it lead to a difference in performance on the (E) RotaRod or (F) Balance Beam tasks (P-Myrf(+/-); n=16, P-Myrf(-/-); n=25) which may have indicated a more general motor skill deficit. (G-H) Sex difference in running speed and activity level. (G) Females (n=57) outperformed males (n=67) on the complex running wheel in terms of running performance (Mixed ANOVA: Main effect of Sex; F(1,120)= 19.98, p<0.001). (H) Females also spent more time active on the wheel over 8 days Mann-Whitney U, p=0.014. (K) Active intervals over 8 days (1 hour time bins) for P-Myrf(-/-) and P-Myrf(+/-) animals. While both Genotypes displayed comparable day/night activity patterns, P-Myrf(-/-) had on average fewer active intervals on the CW during the night when compared to P-Myrf(+/-) (Mixed ANOVA: Main effect of Genotype: (F(1,120)=4.48 p=036). Data presented as Mean ± 95%CI. Data presented as Mean±95%CI or boxplots. Asterix indicate statistical significance (*p<0.05, **p<0.01, ***p<0.001).

**Suppl. Fig. 2.**
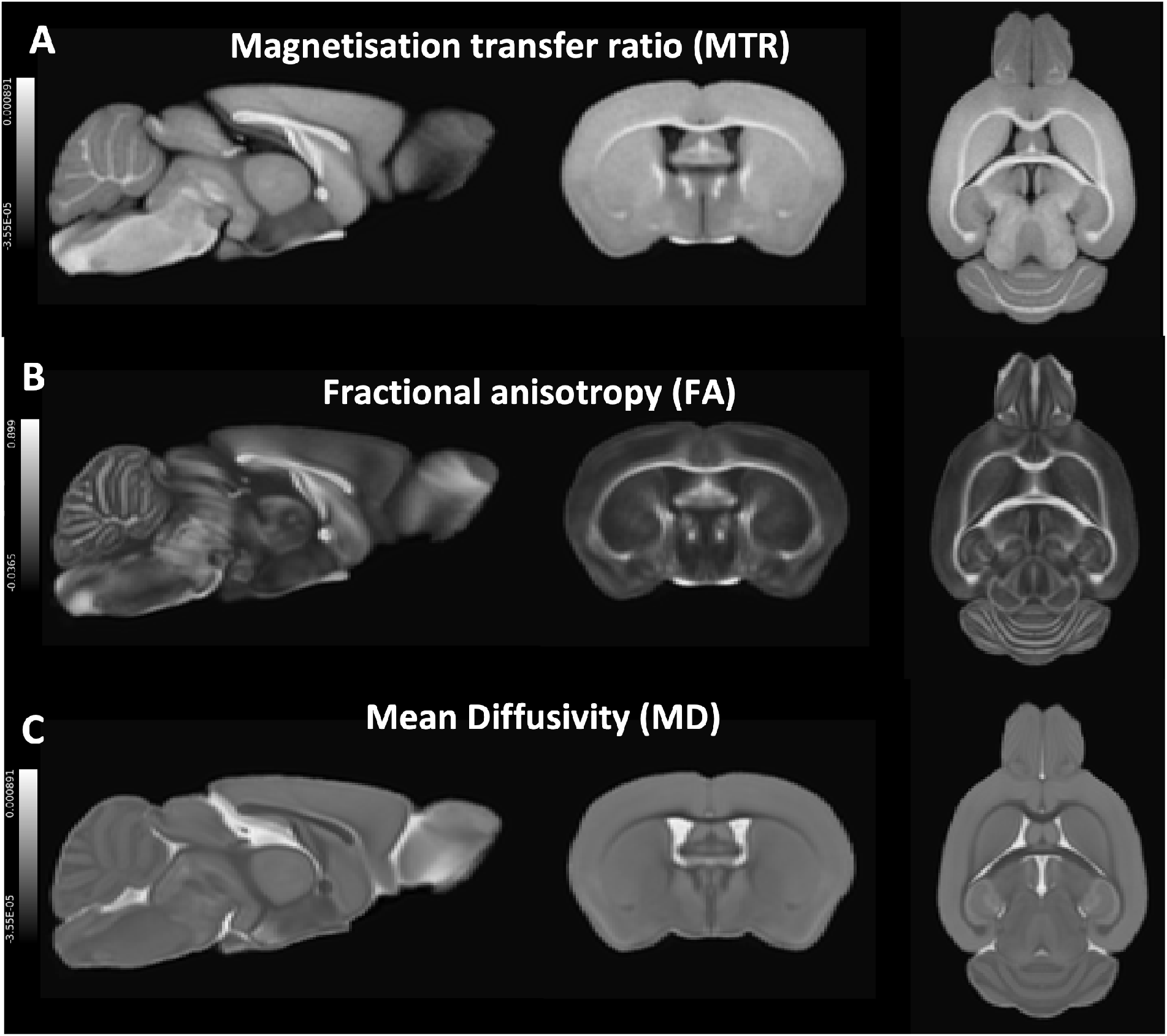
Example images for different parameter contrasts extracted from ex vivo MRI for quantification of brain microstructure. Brain microstructure was assessed using a 9.4 T horizontal bore MR scanner (Varian, Palo Alto, CA, USA) **(A)** Magnetisation transfer ratio (MTR) allows indirect detection of water bound to macromolecules, such as lipids and proteins, and is thus sensitive to myelin (Deloire-Grassin et al. 2000). **(B)** Fractional anisotropy (FA) describes the anisotropy of diffusion of water molecules. **(C)** Mean Diffusivity (MD), which describes the rotationally invariant magnitude of water diffusion within brain tissue. Images shown are the mean of all individual subject images analysed for this study.

**Suppl. Fig. 3.**
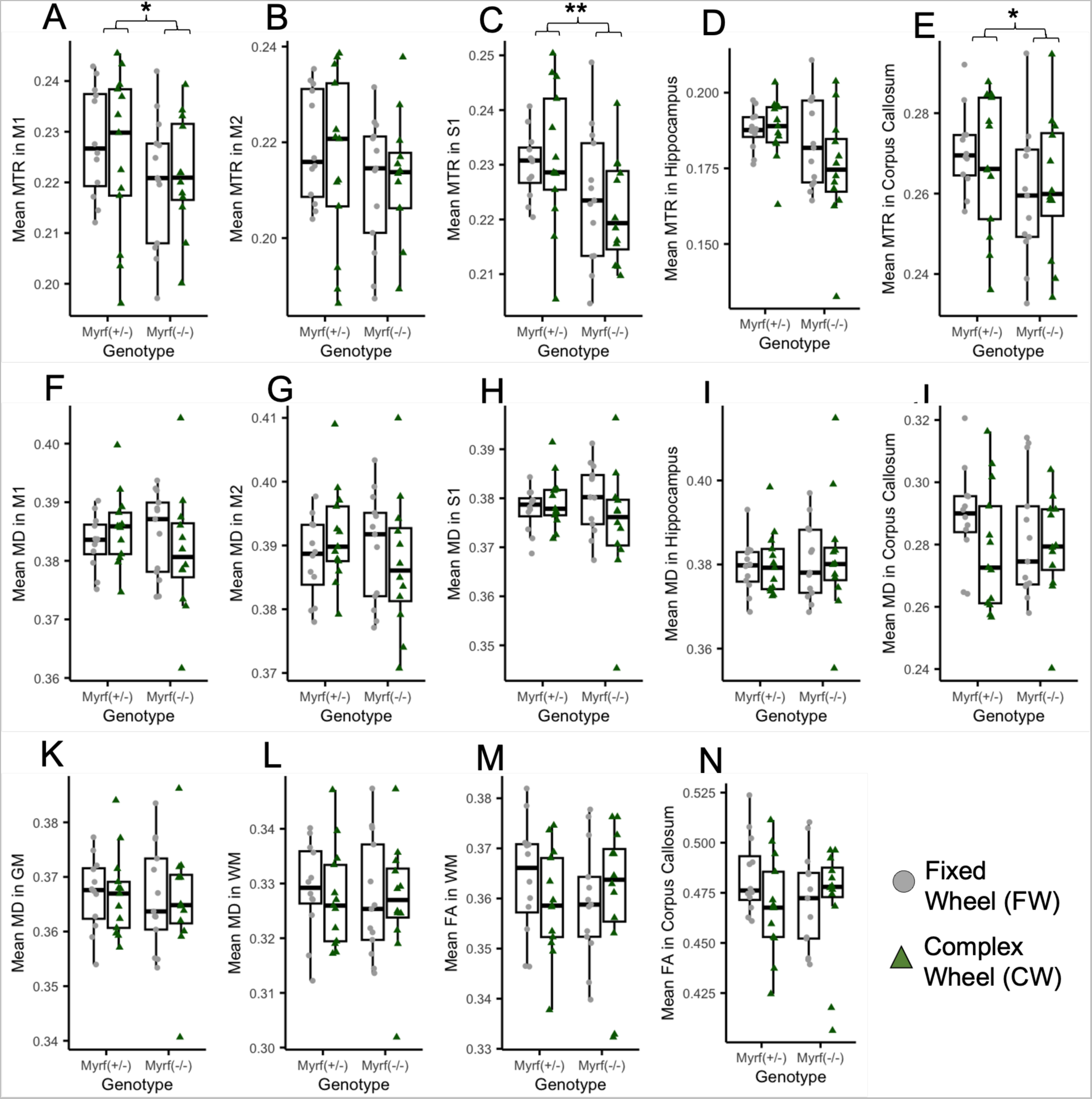
Region of Interest (ROI) Analysis of brain microstructure, as measure by MTR, MD and FA. **(A-E)** MTR values in grey matter ROIs. A Mixed ANOVA revealed a main effect of Genotype in MTR grey matter regions, post-hoc pairwise comparison (Bonferroni adjusted) revealed a significant Genotype effect in the **(A)** Primary motor cortex (M1), **(C)** Somatosensory Cortex (S1), and **(E)** MTR in the anterior Corpus Callosum. **(F-J)** DWI-derived Mean Diffusivity (MD) values in ROI, **(K)** whole brain mask and **(L)** white matter skeleton. **(M-N)** DWI-derived Fractional Anisotropy (FA) values **(M)** white matter skeleton and **(N)** anterior corpus callosum. Asterix indicate statistical significance (*p<0.05, **p<0.01).

**Suppl. Fig. 4.**
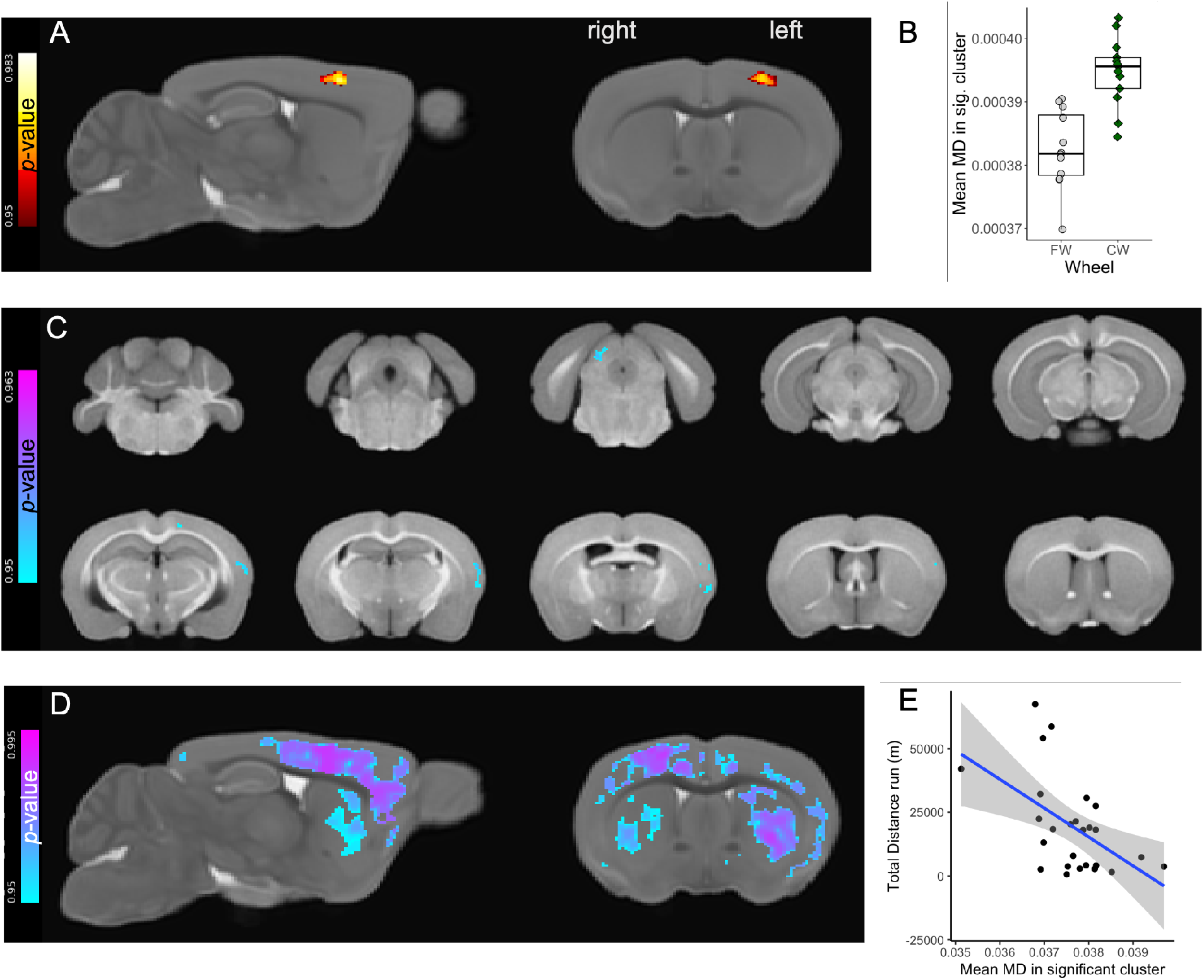
Voxel-wise analysis of DWI-derived parameters revealed an effect wheel running in brain microstructure. Results from Voxel-wise analysis of MRI-derived metrics. (**A**) A cluster in the right primary motor cortex (M1) grey matter region had higher MD for P-Myrf(-/+) control animals that ran on the Complex Wheel (CW), when compared to P-Myrf(-/+) exposed to the fixed wheel (FW). (**B**) Mean MD extracted from cluster depicted in (**A**). **(C)** MTR was lower for P-Myrf(-/-) animals in a few clusters located in cortical regions and the midbrain (superior colliculus). (**D**) MD across a diffuse area across the grey matter was correlated to the total distance ran on the complex running wheel by animals across both genotypes. **(E)** Mean MD extracted from the ROI depicted in **(D)** plotted against the total distance run on the complex wheel. Permutation test (‘randomise’, FSL) used for voxel-wise analysis. Spearman’s test used for correlation analysis.

**Suppl. Fig. 5.**
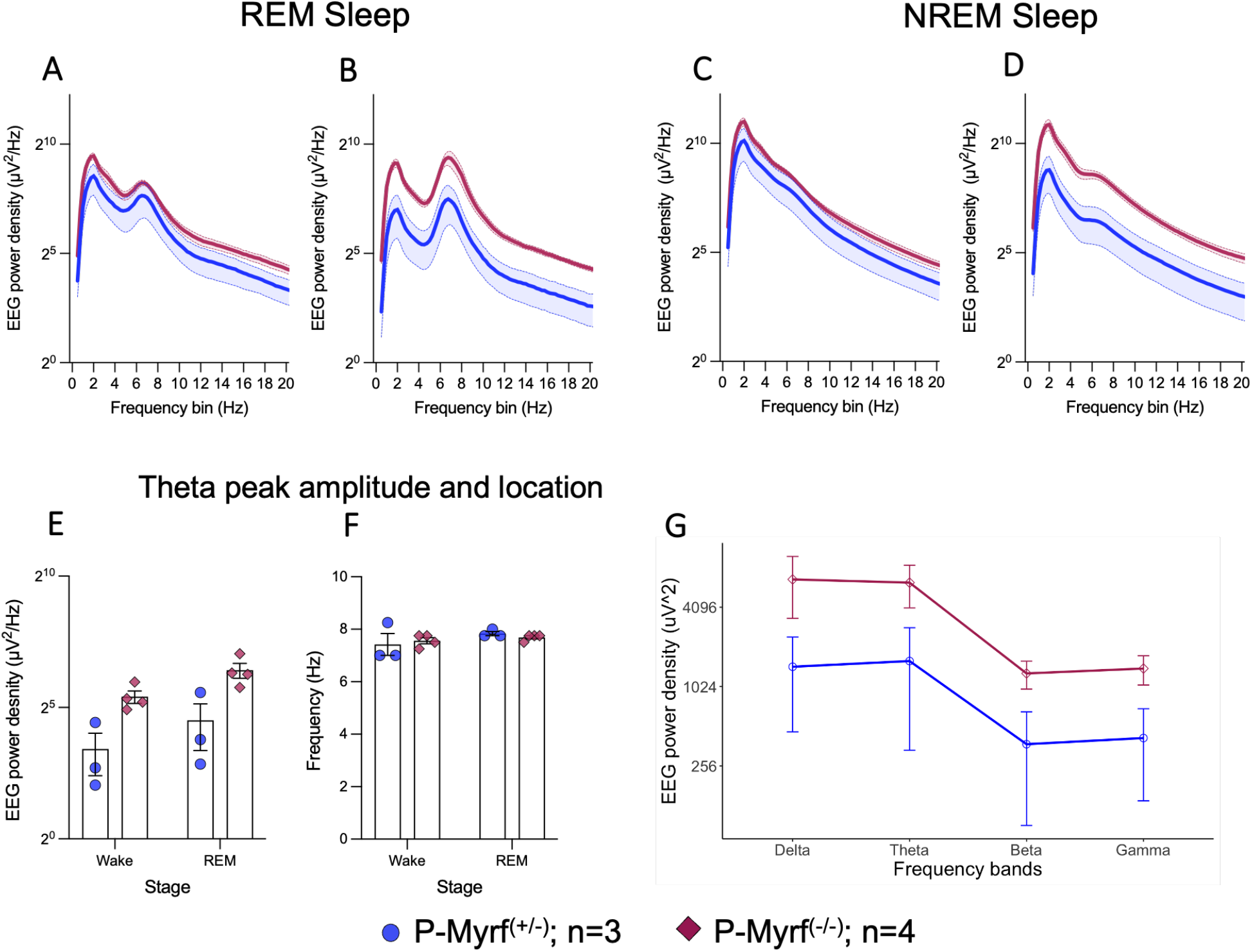
Difference in EEG recording between P-Myrf(-/-) and P-Myrf(+/-). EEG power density by frequency during rapid eye movement (REM) sleep in (**A)** frontal channel and **(B)** occipital channels. EEG power density by frequency during non-REM sleep (NREM) in (**C)** frontal channel and **(D)** occipital channels. **(E)** Amplitude and (**F**) location of peak theta power per vigilance state (Wake, REM) as measured in occipital channel. While peak amplitude was different between the genotype groups, there was no shift in the frequency of the peak. **(G)** Mean group EEG power density (PSD) per frequency bands, delta, δ (0.5–3 Hz), theta, θ (4–12 Hz), beta, β (12.5–25 Hz), and gamma, γ (30-60 Hz). Data presented as Mean±SEM.

**Suppl. Fig. 6.**
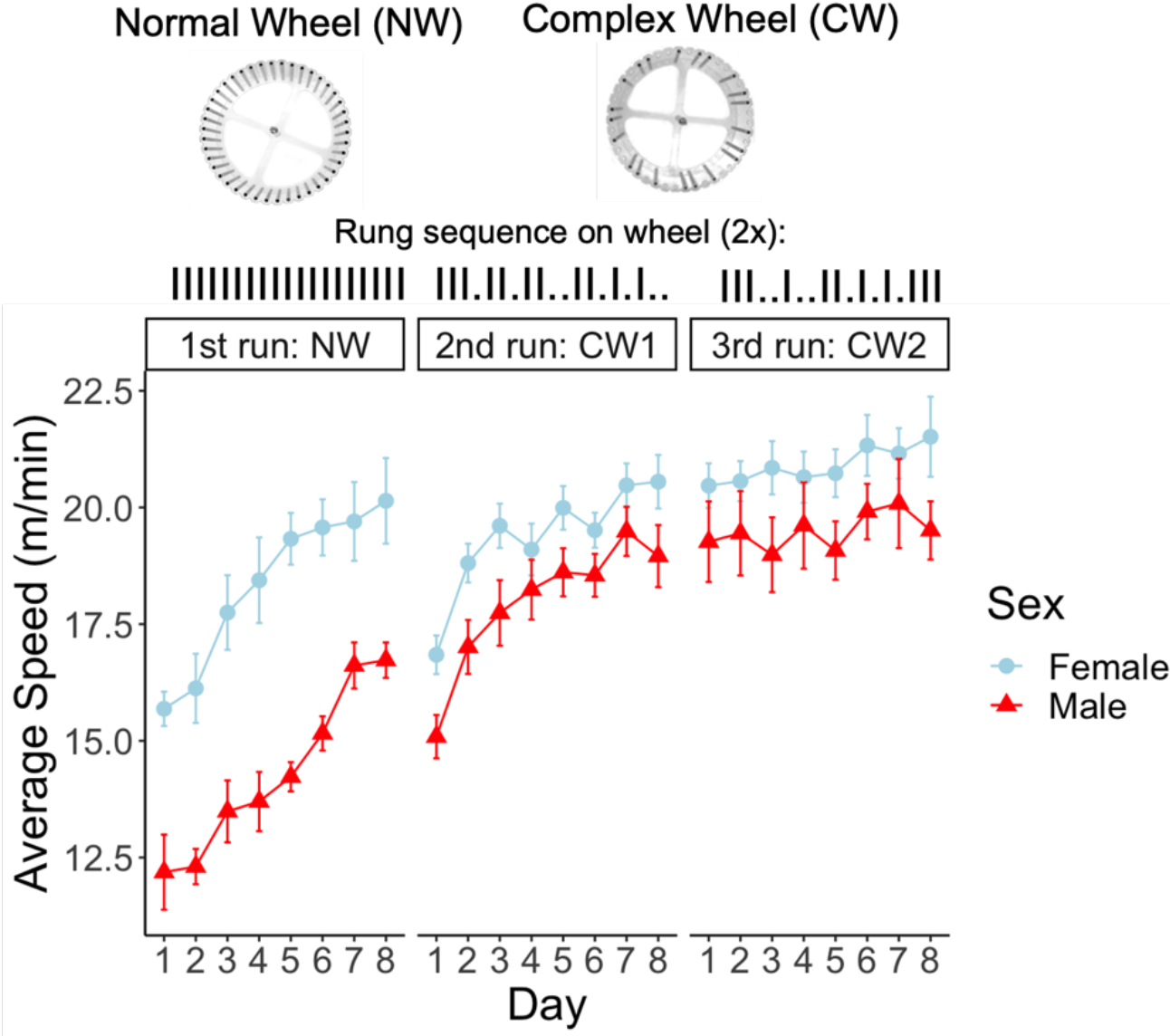
The effect of complex wheel sequence on running performance. Wildtype mice (C58BL/6JOIaHsd, 8 females, 8 males, Age: P84) exposed to a normal running wheel (NW) show significant improvements in their running speed over 8 days (Main effect of Day: F(3.14, 47.1)=21.8, p<0.001), with females outperforming males (Sex*Day Interaction: F(1, 15)=34.6, p<0.001). Switching to complex running wheel (CW1) leads to a temporary reduction in running speed (Day 8 on NW vs Day 1 on CW1; Main effect of wheel: F(1, 14)=27.0, p<0.001), indicating a specific task demand of the complex wheel. However, running speed on day 1 on CW1 is significantly higher than on day 1 on NW (Main effect of wheel: F(1, 14)=12.0, p=0.004), indicating some skill in wheel running was retained between NW and CW1. Switching the rung sequence of the complex running wheel (CW2) after 8 days has no effect on running performance compared to last day of running on CW1. Mixed Two-way ANOVA used for statistical analysis (between-subject factor: Sex; within-subject factor: Time (8 Levels) or Wheel (2 Levels)). Data presented as Mean±SEM.

**Suppl. Fig. 7.**
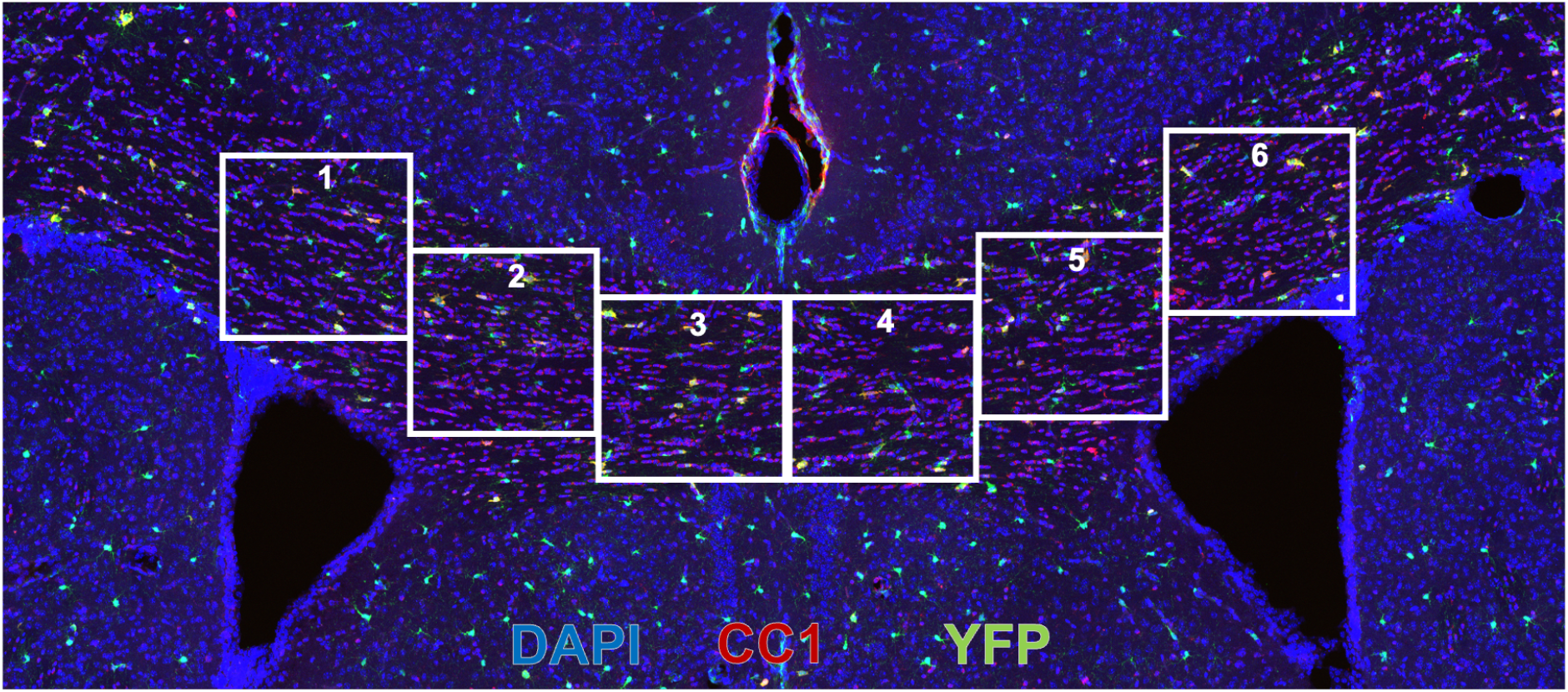
Quantification of oligodendrocyte differentiation in the corpus Callosum. Representative image of a brain slices showing the anterior Corpus Callosum region used for quantification of YFP+ positive OPCs and CC1+ mature Oli-godendrocytes. Six images were taken from each section (three sections per animal), as illustrated by the white frames in the image.

**Suppl. Table 1.**
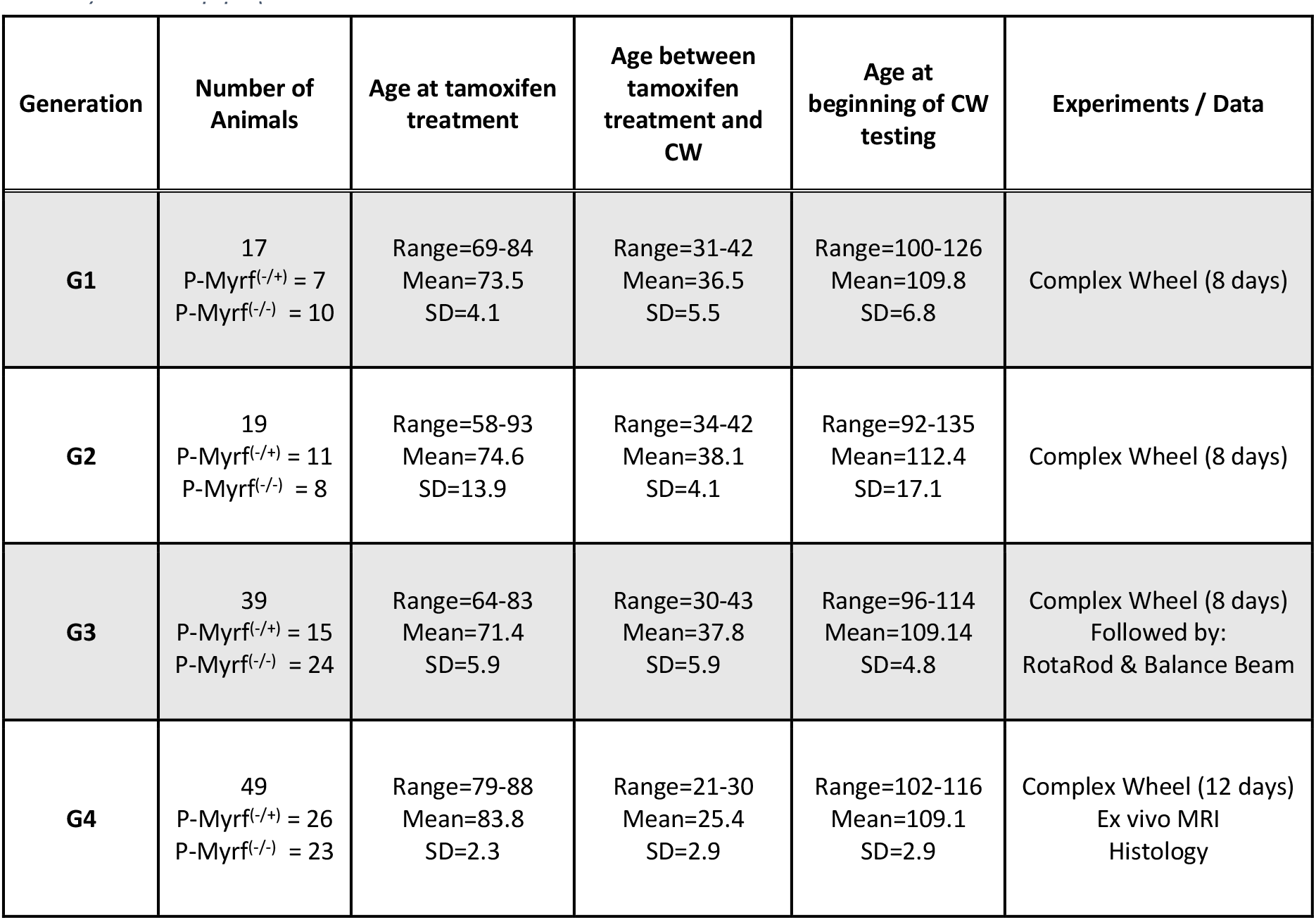
Variability of animal’s age at important experimental timepoints within different generations of Myrf-cKO animals that ran the complex wheel. Age is presented in days after birth

| Generation | Number of Animals | Age at tamoxifen treatment | Age between tamoxifen treatment and CW | Age at beginning of CW testing | Experiments / Data |
| --- | --- | --- | --- | --- | --- |
| **G1** | 17<br>P-Myrf<sup>(-/+)</sup> = 7<br>P-Myrf<sup>(-/-)</sup> = 10 | Range=69-84<br>Mean=73.5<br>SD=4.1 | Range=31-42<br>Mean=36.5<br>SD=5.5 | Range=100-126<br>Mean=109.8<br>SD=6.8 | Complex Wheel (8 days) |
| **G2** | 19<br>P-Myrf<sup>(-/+)</sup> = 11<br>P-Myrf<sup>(-/-)</sup> = 8 | Range=58-93<br>Mean=74.6<br>SD=13.9 | Range=34-42<br>Mean=38.1<br>SD=4.1 | Range=92-135<br>Mean=112.4<br>SD=17.1 | Complex Wheel (8 days) |
| **G3** | 39<br>P-Myrf<sup>(-/+)</sup> = 15<br>P-Myrf<sup>(-/-)</sup> = 24 | Range=64-83<br>Mean=71.4<br>SD=5.9 | Range=30-43<br>Mean=37.8<br>SD=5.9 | Range=96-114<br>Mean=109.14<br>SD=4.8 | Complex Wheel (8 days)<br>Followed by:<br>RotaRod & Balance Beam |
| **G4** | 49<br>P-Myrf<sup>(-/+)</sup> = 26<br>P-Myrf<sup>(-/-)</sup> = 23 | Range=79-88<br>Mean=83.8<br>SD=2.3 | Range=21-30<br>Mean=25.4<br>SD=2.9 | Range=102-116<br>Mean=109.1<br>SD=2.9 | Complex Wheel (12 days)<br>Ex vivo MRI<br>Histology |

**Suppl. Table 2:** Comprehensive information on the statistical analysis methods utilized and the corresponding results for each dataset.

| Figure | Data & Methods | Statistical results |  |  |  |  |
| --- | --- | --- | --- | --- | --- | --- |
| Fig. 1C | Welch's t-test.<br>P-Myrf(+/-), n=12;<br>P-Myrf(-/-), n=15. | YFP+ cells: t = 4.5627, df = 15.462, p-value = 0.000347<br>YFP+CC1+: t = 7.1621, df = 11.274, p-value = 1.61e-05<br>YFP+CC1-: t = 0.72468, df = 24.246, p-value = 0.4756 |  |  |  |  |
| Fig. 1D | Welch's t-test.<br>P-Myrf(+/-), n=12;<br>P-Myrf(-/-), n=15. | YFP+CC1+/YFP+CC1-: t = 10.199, df = 11.338, p-value = 4.706e-07 |  |  |  |  |
| Fig. 2D | Mixed Two-way ANOVA<br>P-Myrf(+/-), n=59<br>(Female, n=28);<br>P-Myrf(-/-), n=65<br>(Female, n=29);<br>Between subject factor:<br>MYRFGenotype (2 levels),<br>Sex (2 levels)<br>Within subject factor:<br>Time (Days, 8 levels) | Effect | DFn | DFd | F | p |
|  |  | (Intercept) | 1 | 120 | 900.615 | <0.001 |
|  |  | MYRFGenotype | 1 | 120 | 10 | 0.002 |
|  |  | Sex | 1 | 120 | 13.081 | <0.001 |
|  |  | Time | 2.74 | 328.53 | 170.149 | <0.001 |
|  |  | MYRFGenotype:Sex | 1 | 120 | 3.041 | 0.084 |
|  |  | MYRFGenotype:Time | 2.74 | 328.53 | 5.569 | 0.001 |
|  |  | Sex:Time | 2.74 | 328.53 | 6.344 | <0.001 |
| Fig. 2E | Mixed Two-way ANOVA<br>P-Myrf(+/-), n=59<br>(Female, n=28);<br>P-Myrf(-/-), n=65<br>(Female, n=29);<br>Between subject factor:<br>MYRFGenotype (2 levels),<br>Sex (2 levels)<br>Within subject factor:<br>Time (Days, 8 levels) | Effect | DFn | DFd | F | p |
|  |  | (Intercept) | 1 | 120 | 1791.129 | <0.001 |
|  |  | MYRFGenotype | 1 | 120 | 11.922 | <0.001 |
|  |  | Sex | 1 | 120 | 17.481 | <0.001 |
|  |  | Time | 2.57 | 308.96 | 185.786 | <0.001 |
|  |  | MYRFGenotype:Sex | 1 | 120 | 1.867 | 0.174 |
|  |  | MYRFGenotype:Time | 2.57 | 308.96 | 4.764 | 0.005 |
|  |  | Sex:Time | 2.57 | 308.96 | 4.614 | 0.006 |
| Fig. 2F | Mann-Whitney U test<br>P-Myrf(+/-), n=59<br>(Female, n=28);<br>P-Myrf(-/-), n=65<br>(Female, n=29); | W = 2504, p-value = 0.003369 |  |  |  |  |
| Fig. 2G | Mann-Whitney U test<br>P-Myrf(+/-), n=59<br>(Female, n=28);<br>P-Myrf(-/-), n=65<br>(Female, n=29); | W = 2536, p-value = 0.001988 |  |  |  |  |
| Fig. 2H | Mixed Two-way ANOVA<br>P-Myrf(+/-), n=59<br>(Female, n=28);<br>P-Myrf(-/-), n=65<br>(Female, n=29);<br>Between subject factor:<br>MYRFGenotype (2 levels),<br>Sex (2 levels)<br>Within subject factor:<br>Time (12 levels (20 mins intervals)) | Effect | DFn | DFd | F | p |
|  |  | MYRFGenotype | 1 | 120 | 2.727 | 0.101 |
|  |  | Sex | 1 | 120 | 0.753 | 0.387 |
|  |  | Time | 4.83 | 579.63 | 62.791 | <0.001 |
|  |  | MYRFGenotype:Sex | 1 | 120 | 5.446 | 0.021 |
|  |  | MYRFGenotype:Time | 4.83 | 579.63 | 2.627 | 0.025 |
|  |  | Sex:Time | 4.83 | 579.63 | 3.088 | 0.01 |
| Fig. 2I | Mann-Whitney U test<br>P-Myrf(+/-), n=59<br>(Female, n=28);<br>P-Myrf(-/-), n=65<br>(Female, n=29); | W = 2413.5, p-value = 0.01317 |  |  |  |  |
| Fig. 2J | Mann-Whitney U test<br>P-Myrf(+/-), n=59<br>(Female, n=28);<br>P-Myrf(-/-), n=65<br>(Female, n=29); | W = 2411, p-value = 0.01364 |  |  |  |  |
| Fig. 2K | Spearman's rank correlation rho: | Across Genotype: S = 63551, p-value <0.001, rho = 0.7999962<br>P-Myrf(+/-): S = 12760, p-value <0.001, rho 0.6271278<br>P-Myrf(-/-): S = 5874.6, p-value <0.001, rho 0.8716223 |  |  |  |  |
|  | cor.test(a, b<br>method=c("spearman"))<br>P-Myrf(+/-), n=59<br>(Female, n=28);<br>P-Myrf(-/-), n=65<br>(Female, n=29); |  |  |  |  |  |
| Fig. 2L | Mixed Two-way ANOVA<br>P-Myrf(+/-), n=59<br>(Female, n=28);<br>P-Myrf(-/-), n=64<br>(Female, n=29);<br>Between subject factor:<br>MYRFGenotype (2<br>Levels), Sex (2 Levels)<br>Within subject factor:<br>Time (6 levels (10 mins<br>intervals)) | Effect | DFn | DFd | F | p |
|  |  | MYRFGenotype | 1 | 120 | 2.848 | 0.094 |
|  |  | Sex | 1 | 120 | 0.058 | 0.81 |
|  |  | Time | 2.21 | 265.32 | 88.001 | <0.001 |
|  |  | MYRFGenotype:Sex | 1 | 120 | 2.846 | 0.094 |
|  |  | MYRFGenotype:Time | 2.21 | 265.32 | 9.06 | <0.001 |
|  |  | Sex:Time | 2.21 | 265.32 | 1.81 | 0.161 |
| Fig. 2M | Mann-Whitney U test<br>P-Myrf(+/-), n=59<br>(Female, n=28);<br>P-Myrf(-/-), n=65<br>(Female, n=29); | W = 2490, p-value = 0.00421 |  |  |  |  |
| Fig. 2N | Mixed Two-way ANOVA<br>P-Myrf(+/-), n=59<br>(Female, n=28);<br>P-Myrf(-/-), n=64<br>(Female, n=29);<br>Between subject factor:<br>MYRFGenotype (2<br>Levels), Sex (2 Levels)<br>Within subject factor:<br>Time (5 Intervals (1H<br>intervals)) | Effect | DFn | DFd | F | p |
|  |  | MYRFGenotype | 1 | 118 | 4.998 | 0.027 |
|  |  | Sex | 1 | 118 | 3.09 | 0.081 |
|  |  | Time | 2.2 | 259.93 | 53.822 | <0.001 |
|  |  | MYRFGenotype:Sex | 1 | 118 | 3.709 | 0.057 |
|  |  | MYRFGenotype:Time | 2.2 | 259.93 | 1.224 | 0.298 |
|  |  | Sex:Time | 2.2 | 259.93 | 3.045 | 0.044 |
| Fig. 2O | Mann-Whitney U test<br>P-Myrf(+/-), n=58<br>(Female, n=28);<br>P-Myrf(-/-), n=64<br>(Female, n=29); | W = 2278, p-value = 0.0307 |  |  |  |  |
| Fig. 3D | Mixed Two-way ANOVA<br>P-Myrf(+/-), n=12<br>P-Myrf(-/-), n=13<br>Between subject factor:<br>MYRFGenotype (2<br>Levels), Batch (4 Levels)<br>Within subject factor:<br>ROI (4 Levels) | Effect | DFn | DFd | F | p |
|  |  | MYRFGenotype | 1 | 17 | 7.318 | 0.015 |
|  |  | Batch | 3 | 17 | 0.889 | 0.467 |
|  |  | ROI | 1.27 | 21.57 | 190.596 | <0.001 |
|  |  | MYRFGenotype:Batch | 3 | 17 | 2.786 | 0.072 |
|  |  | MYRFGenotype:ROI | 1.27 | 21.57 | 2.084 | 0.161 |
|  |  | Batch:ROI | 3.81 | 21.57 | 3.034 | 0.041 |
| Fig. 3D | Two-way ANOVAs<br>P-Myrf(+/-), n=12<br>P-Myrf(-/-), n=13<br>Between subject factor:<br>MYRFGenotype (2<br>Levels), Batch (4 Levels) | ROI | Effect | F | p | p (adjusted) |
|  |  | MTRHipp | MYRFGenotype | 0.969 | 0.339 | 1 |
|  |  | MTRM1 | MYRFGenotype | 10.523 | 0.005 | 0.02 |
|  |  | MTRM2 | MYRFGenotype | 9.029 | 0.008 | 0.032 |
|  |  | MTRS1 | MYRFGenotype | 5.224 | 0.035 | 0.14 |
| Fig. 3E | Two-way ANOVA<br>P-Myrf(+/-), n=12<br>P-Myrf(-/-), n=13<br>Between subject factor:<br>MYRFGenotype (2<br>Levels), Batch (4 Levels) | ROI | DFn | DFd | F | p |
|  |  | MTRaCCWMSkel | 1 | 17 | 7.77 | 0.013 |
| Fig. 3G | Grey matter skeleton -<br>MTR<br>Two-way ANOVA<br>P-Myrf(+/-), n=25<br>P-Myrf(-/-), n=25<br>Between subject factor:<br>MYRFGenotype (2<br>Levels), Batch (4 Levels) | Effect | DFn | DFd | F | p |
|  |  | MYRFGenotype | 1 | 34 | 6.04 | 0.019 |
|  |  | Wheel | 1 | 34 | 0.065 | 0.8 |
|  |  | Batch | 3 | 34 | 2.179 | 0.108 |
|  |  | MYRFGenotype:Wheel | 1 | 34 | 0.061 | 0.807 |
|  |  | MYRFGenotype:Batch | 3 | 34 | 1.117 | 0.356 |
|  |  | Wheel:Batch | 3 | 34 | 0.395 | 0.758 |
| Fig. 3H | White matter skeleton - MTR<br>Two-way ANOVA<br>P-Myrf(+/-), n=25<br>P-Myrf(-/-), n=25<br>Between subject factor: MYRFGenotype (2 Levels),<br>Wheel (2 Levels), Batch (4 Levels) | **Effect** | **DFn** | **DFd** | **F** | **p** |
|  |  | MYRFGenotype | 1 | 34 | 4.728 | 0.037 |
|  |  | Wheel | 1 | 34 | 0.028 | 0.869 |
|  |  | Batch | 3 | 34 | 1.563 | 0.216 |
|  |  | MYRFGenotype:Wheel | 1 | 34 | 0.083 | 0.775 |
|  |  | MYRFGenotype:Batch | 3 | 34 | 0.939 | 0.433 |
|  |  | Wheel:Batch | 3 | 34 | 0.697 | 0.56 |
| Fig. 4E | Mixed Two-way ANOVA<br>P-Myrf(+/-), n=3<br>P-Myrf(-/-), n=4<br>Between subject factor: MYRFGenotype (2 Levels),<br>Within subject factor: EEG location (2 Levels) | **Effect** | **DFn** | **DFd** | **F** | **p** |
|  |  | Genotype | 1 | 5 | 10.398 | 0.023 |
|  |  | EEG location | 1 | 5 | 4.207 | 0.096 |
|  |  | Genotype: EEG location | 1 | 5 | 0.779 | 0.418 |
| Fig. 4F | Mixed Two-way ANOVA<br>P-Myrf(+/-), n=3<br>P-Myrf(-/-), n=4<br>Between subject factor: MYRFGenotype (2 Levels),<br>Within subject factor: EEG location (2 Levels) | **Effect** | **DFn** | **DFd** | **F** | **p** |
|  |  | Genotype | 1 | 5 | 12.751 | 0.016 |
|  |  | EEG location | 1 | 5 | 0.155 | 0.71 |
|  |  | Genotype: EEG location | 1 | 5 | 1.463 | 0.28 |
| Fig. 4H | Mixed Two-way ANOVA<br>P-Myrf(+/-), n=3<br>P-Myrf(-/-), n=4<br>Between subject factor: MYRFGenotype (2 Levels),<br>Within subject factor: EEG location (2 Levels) | **Effect** | **DFn** | **DFd** | **F** | **p** |
|  |  | Genotype | 1 | 5 | 11.385 | 0.02 |
|  |  | EEG location | 1 | 5 | 1.762 | 0.242 |
|  |  | Genotype:EEG location | 1 | 5 | 1.071 | 0.348 |
| Suppl. Fig. 1H | Mann–Whitney U test<br>Males: n=67;<br>Females: n=57; | W = 2411, p-value = 0.01364 |  |  |  |  |
| Suppl. Fig 1I | Mixed Two-way ANOVA<br>P-Myrf(+/-), n=59 (Female, n=28);<br>P-Myrf(-/-), n=64 (Female, n=29);<br>Between subject factor: MYRFGenotype (2 Levels), Sex (2 Levels)<br>Within subject factor: Time (192 Intervals – 1h size)) | **Effect** | **DFn** | **DFd** | **F** | **p** |
|  |  | MYRFGenotype | 1 | 120 | 4.484 | 0.036 |
|  |  | Sex | 1 | 120 | 30.276 | <0.001 |
|  |  | Interval | 191 | 22920 | 103.115 | <0.001 |
|  |  | MYRFGenotype:Sex | 1 | 120 | 0.604 | 0.438 |
|  |  | MYRFGenotype:Interval | 191 | 22920 | 2.12 | <0.001 |
|  |  | Sex:Interval | 191 | 22920 | 8.859 | <0.001 |
| Suppl. Fig 3A-D | Mixed Two-way ANOVA<br>P-Myrf(+/-), n=25<br>P-Myrf(-/-), n=25<br>Between subject factor: MYRFGenotype (2 Levels), Batch (4 Levels)<br>Within subject factor: ROI (4 Levels) | **Effect** | **DFn** | **DFd** | **F** | **p** |
|  |  | MYRFGenotype | 1 | 34 | 10.464 | 0.003 |
|  |  | Wheel | 1 | 34 | 0.196 | 0.661 |
|  |  | Batch | 3 | 34 | 3.426 | 0.028 |
|  |  | ROI_GM_MTR | 1.27 | 43.21 | 299.344 | <0.001 |
|  |  | MYRFGenotype:Wheel | 1 | 34 | 0.076 | 0.784 |
|  |  | MYRFGenotype:Batch | 3 | 34 | 2.39 | 0.086 |
|  |  | Wheel:Batch | 3 | 34 | 0.455 | 0.715 |
|  |  | MYRFGenotype:ROI_GM_MTR | 1.27 | 43.21 | 0.196 | 0.719 |
|  |  | Wheel:ROI_GM_MTR | 1.27 | 43.21 | 0.34 | 0.615 |
| Suppl. Fig 3A-D | Two-way ANOVAs<br>P-Myrf(+/-), n=25<br>P-Myrf(-/-), n=25 | **ROI** | **Effect** | **F** | **p** | **p (adjusted)** |
|  |  | MTRHipp | MYRFGenotype | 6.142 | 0.018 | 0.072 |
|  |  | MTRM1 | MYRFGenotype | 7.814 | 0.008 | 0.032 |
|  |  | MTRM2 | MYRFGenotype | 4.816 | 0.035 | 0.14 |
|  | Between subject factor:<br>MYRFGenotype (2 Levels), Batch (4 Levels) | MTRS1 | MYRFGenotype | 13.376 | <0.001 | 0.0032 |
| Suppl.<br>Fig.<br>3E | White matter skeleton -<br>MTR<br>Two-way ANOVA<br>P-Myrf(+/-), n=25<br>P-Myrf(-/-), n=25<br>Between subject factor:<br>MYRFGenotype (2 Levels),<br>Wheel (2 Levels), Batch (4 Levels) | **Effect** | **DFn** | **DFd** | **F** | **p** |
|  |  | MYRFGenotype | 1 | 34 | 6.574 | 0.015 |
|  |  | Wheel | 1 | 34 | 0.1 | 0.754 |
|  |  | Batch | 3 | 34 | 4.91 | 0.006 |
|  |  | MYRFGenotype:Wheel | 1 | 34 | 0.621 | 0.436 |

